# Cell intercalation driven by SMAD3 underlies secondary neural tube formation

**DOI:** 10.1101/2020.08.24.261008

**Authors:** Elena Gonzalez-Gobartt, José Blanco-Ameijeiras, Susana Usieto, Guillaume Allio, Bertrand Benazeraf, Elisa Martí

## Abstract

Body axis elongation is a hallmark of the vertebrate embryo, involving the architectural remodelling of the tailbud. Although it is clear how bi-potential neuro-mesodermal progenitors (NMPs) contribute to embryo elongation, the dynamic events that lead to *de novo* lumen formation and that culminate in the formation of a 3-Dimensional, secondary neural tube from NMPs, are poorly understood. Here, we used in vivo imaging of the chicken embryo to show that cell intercalation downstream of TGF-beta/SMAD3 signalling is required for secondary neural tube formation. Our analysis describes the initial events in embryo elongation including lineage restriction, the epithelial-to-mesenchymal transition of NMPs, and the initiation of lumen formation. Importantly, we show that the resolution of a single, centrally positioned continuous lumen, which occurs through the intercalation of central cells, requires SMAD3 activity. We anticipate that these findings will be relevant to understand caudal, skin-covered neural tube defects, amongst the most frequent birth defects detected in humans.

**HIGHLIGHTS:**

.**- Initiation of the lumen formation follows the acquisition of neural identity and epithelial polarization**.
.**- Programmed cell death is not required for lumen resolution**.
.**- Resolution of a single central lumen requires cell intercalation, driven by Smad3 activity**
.**- The outcome of central cell division preceding cell intercalation, varies along the cranio-caudal axis**.

## INTRODUCTION

A proliferative cell population at the posterior end of the vertebrate embryo, known as neuromesodermal progenitors (NMPs), sustained body axis elongation in vertebrates (Cambray & Wilson, 2007; Henrique et al., 2015; Tzouanacou et al., 2009). These cells have been found in zebrafish, chick, mouse and human embryos, and they produce both the neural tissue that makes-up the caudal spinal cord, and mesodermal tissues like muscle and bone (Cambray & Wilson, 2007; Henrique et al., 2015; Kölliker, 1884; McGrew et al., 2008). NMPs have attracted much attention of late having established protocols to generate them *in vitro* from human pluripotent stem cells (Gouti et al., 2014). This advance has not only allowed the regulatory gene networks that determined NMP fate specification and lineage restriction to be deciphered (Gouti et al., 2017; Metzis et al., 2018), but it has also provided a new experimental paradigm to study the cellular and molecular basis of caudal spinal cord generation. However, despite their relevance to understand caudal skin-covered neural tube defects (NTDs: (Greene & Copp, 2014; Morris et al., 2016; Saitsu & Shiota, 2008), the signaling pathways and cellular events required to shape the secondary neural tube (SNT) from NMPs in 3D are poorly understood.

Morphogenesis of the SNT involves the mesenchymal-to-epithelial transition (MET) of NMPs, which is concomitant to the lineage restriction and the confinement of these cells within a solid medullary cord, surrounded by a growing basement membrane. The subsequent cavitation of this chord is required to form the SNT (Catala et al., 1995; Colas & Schoenwolf, 2001; Griffith et al., 1992; Shimokita & Takahashi, 2011). The limited availability of human tissue to perform histological analyses at different developmental stages (Abbott, 2011) reinforces the need to use animal models to understand the events that shape the SNT, particularly since NTDs are among the most common human birth defects (Greene & Copp, 2014). Here, we have used the chick embryo to examine the signalling proteins that control the subcellular events and the 3D tissue rearrangements shaping the SNT. At early stages of development, the avian body plan is very similar to that of mammals, and the thin and planar nature of chick embryos endows them with excellent optical properties (Le Douarin et al., 1998; Rupp et al., 2003; Shimokita & Takahashi, 2011). Furthermore, anatomically the SNT of the chick embryo extends up to the lumbar region (Criley, 1969; Dady et al., 2014), closely resembling human development (O’Rahilly & Muller, 1994, 2002; Saitsu et al., 2004; Saitsu & Shiota, 2008), whereas SNT only occurs at the level of the tail in mice embryos (Nievelstein et al., 1993; Schoenwolf, 1984; Shum et al., 2010).

Here we found Transforming Growth Factor-beta (TGF-β) signalling to be active during SNT formation. Members of the TGF-β superfamily have been implicated as major instructive signals of the epithelial-to-mesenchymal transition (EMT) in many morphogenetic processes, including gastrulation and mesendodermal ingression from the embryonic epiblast (Luxardi et al., 2010), and in the delamination of neural crest cells from the developing neural tube (Theveneau & Mayor, 2012). From loss-of-function experiments in the chick embryo, we show here that the initial events in SNT formation appear to be independent of SMAD3 signalling, involving the lineage restriction of dual fated NMPs, as well as the MET and full epithelialization of caudal neural progenitor cells. However, establishing a single, centrally positioned continuous lumen occurs through the intercalation of central cells and this does appear to require SMAD3 activity. Together, we describe here a novel activity for TGF-β/SMAD3 that we believe may be relevant to the development of human caudal skin-covered NTDs.

## RESULTS

### Secondary neural tube formation in the chick embryo requires TGF-β mediated Smad3 activity

During vertebrate development, NMPs are recruited to elongate the caudal body axis and to drive the elongation of the neural tube (NT). These bi-potential cells will generate neural progenitor cells (NPCs) of the caudal medullary cord by undergoing a MET, these forming the SNT. The process is completed by the cavitation of the medullary cord and it can be followed along the cranio-caudal axis of a HH15 chick embryo (Hamburger & Hamilton, 1951), 50-55 hours post fertilisation (Fig. 1A-E). The first cells to epithelialize are confined to the periphery of the medullary chord and then surrounded by a developing basement membrane, while the central cells remain mesenchymal until the very end of the process. Small cavities of varied size form between these two cell populations, later resolving to form a single central lumen (Fig. 1A’-E’: (Schoenwolf & Delongo, 1980; Schoenwolf & Kelley, 1980).

**FIGURE 1:**
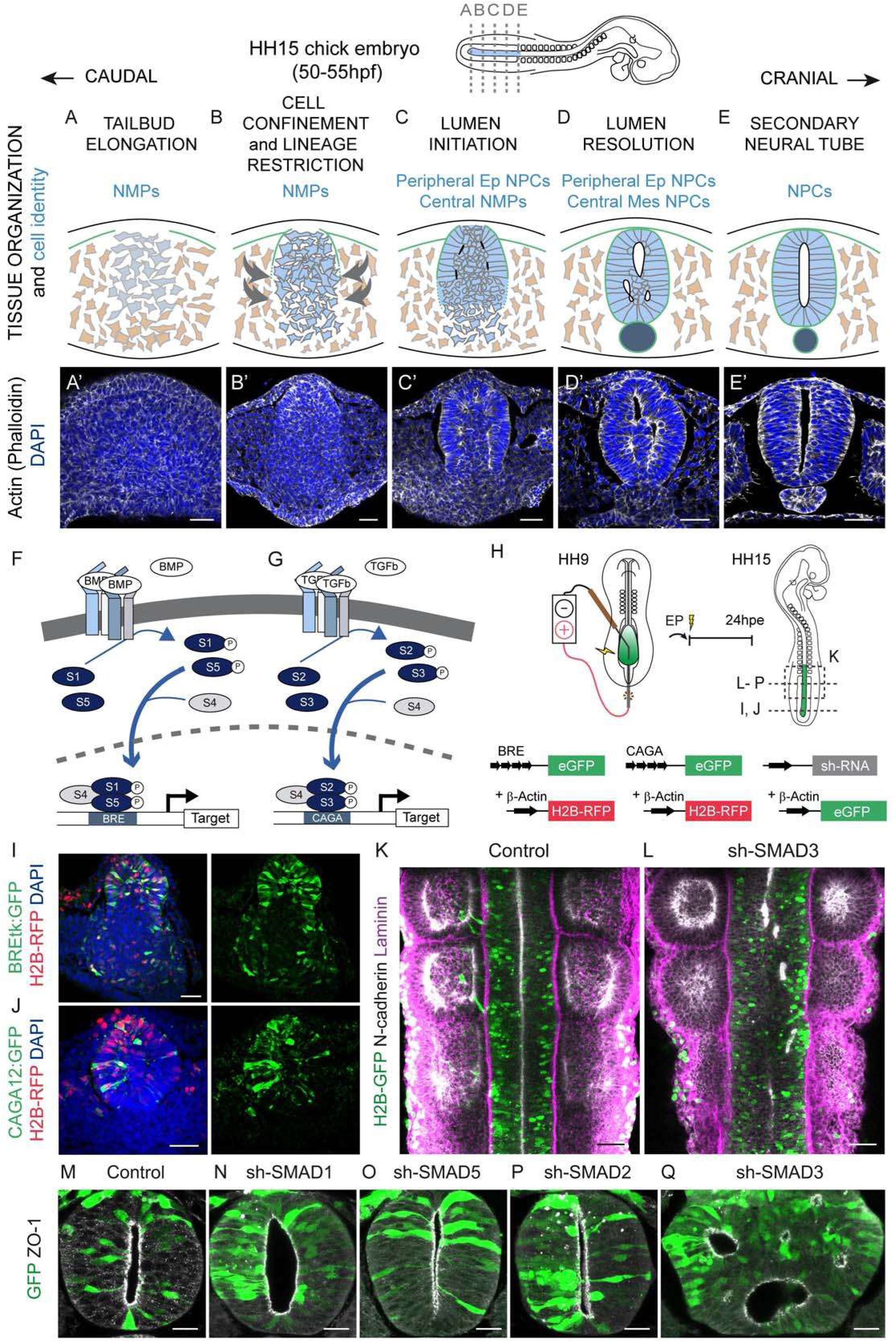
SNT formation in the chick embryos requires TGFβ mediated SMAD3 activity. **(A-E)** Scheme showing a stage HH15 chick embryo, and the cellular processes occurring along the posterior to anterior axis during secondary neural tube (SNT) formation. SNT cells are shown in light blue, the surrounding mesoderm is in brown and the notochord appears in dark blue. **(A’-E’)** Transverse sections at different cranio-caudal levels showing the distribution of actin (white), with DAPI (blue) staining the cell nuclei. Scale bars = 40 μm. **(F**,**G)** Scheme summarizing the BMP and TGF-β signalling components. **(H)** Scheme showing the method and timing of chick embryo electroporation, and the DNA constructs used: EP, electroporation; hpe, hours post-electroporation. **(I)** The BMP reporter is active in NMPs 24 hpe of BRE:GFP (green) and control H2B-RFP (red). DAPI (blue) stains the nuclei. Scale bar = 40 μm. **(J)** The TGF-β reporter is active in NMPs 24 hpe of CAGA12:GFP (green) and control H2B-RFP (red). DAPI (blue) stains the nuclei. Scale bar = 40 μm. **(K**,**L)** Dorsal views of control H2B-GFP (green, F) or sh-SMAD3 (green, G) electroporated NTs. N-cadherin (white) lines the NT lumen and laminin (pink) stains the basement membrane and somites. Scale bars = 40 μm. **(M-Q)** Selected images of transverse sections 24 hpe of the indicated DNAs (green), ZO-1 staining (white) lines the NT lumen. Electroporation of sh-SMAD3 results in multiple lumens (L). Scale bar = 20 μm.

Members of the Wnt and FGF signaling pathways play relevant roles in the control of NMP growth and maintenance, and their differentiation into neural or mesodermal lineages (Turner et al., 2014) although very little is known about signals that instruct the 3D shaping of the caudal neural tube. Here we analysed the expression of ligands of the main developmental pathways in the tail bud of stage HH15 chick embryos. Whole mount in situ hybridization and transverse sections through these same embryos revealed that the expression of BMP2 and TGF-β1 may potentially be associated with SNT formation (Supp. Fig. S1A-D). Interestingly, the TGF-β/BMP pathways are known to play key roles in anterior NT formation (reviewed in Le Dreau & Marti, 2012; Ulloa & Briscoe, 2007), yet their roles in SNT formation are less well understood. Canonical TGF-β/BMP signaling is in general linear, with ligands binding to a defined receptor complex composed of two transmembrane serine/threonine kinases that in turn propagate the signal through the SMAD family of transcription factors (Fig. 1F,G). In situ hybridization experiments to study the distribution of SMAD mRNA, revealed strong SMAD3 expression in polarizing NMPs at the dorsal periphery of the chord, later spreading to the entire developing SNT (Supp. Fig. S2A-D).

We next assessed the endogenous activity of the TGF-β/BMP pathways *in vivo* by electroporating a BMP (BRE:eGFP: Le Dreau et al., 2014; Le Dreau et al., 2012) or a TGF-β responsive (CAGA12-GFP; (Miguez et al., 2013) fluorescent reporter construct, together with a control H2B-RFP vector (Fig. 1H). Both the TGF-β and BMP pathways were active in the elongating chick embryo tail bud at 24 hours post-electroporation (hpe: Fig. 1I,J; see also Supp. Fig. S2J). Electroporation of the TGF-β-responsive CAGA12:GFP fluorescent reporter that is specific to SMAD3 activity (Miguez et al., 2013), together with the control H2B-RFP vector, showed strong activity in the caudal and SNT region at 24 hpe, whereas control H2B-RFP+ cells were found along the whole cranio-caudal axis (Supp. Fig. S2J).

To test the contribution of canonical TGF-β/BMP signalling to SNT formation, we analysed the consequence of SMAD inhibition through the electroporation of short-hairpin (sh)RNA targeting specific chick *SMAD* transcripts. BMP signalling appeared to be dispensable for the correct morphogenesis of the SNT, since inhibition of SMAD1/5 did not impede the formation of a normal SNT (Fig. 1M-O). Similarly, while electroporation of sh-SMAD2 efficiently inhibited endogenous SMAD2 expression (Miguez et al., 2013), it was insufficient to perturb SNT formation (Fig. 1P). However, inhibition of the TGF-β effector SMAD3 produced an aberrant SNT that contained multiple small lumens distributed along the cranio-caudal (Fig. 1K,L) and the dorso-ventral axes (Fig. 1Q). While sh-SMAD3 electroporation efficiently diminished the endogenous SMAD3 (median ±IQR integrated density control = 543.1 ±486.7 vs sh-SMAD3 = 148.6 ±113.5: Supp. Fig. S2K,L), it did not compromise either tissue or NMP viability, as assessed by the rate of apoptosis and of cell proliferation of the SMAD3 deficient (mean ±SD %c-Casp3^+^ cells control = 1.9 ±1.0 vs sh-SMAD3 = 3.0 ±1.8: Supp. Fig S2N; mean ±SD %pH3^+^ cells control = 4.7 ±1.5 vs sh-SMAD3 =3.8 ±1.1: Supp. Fig S2O), or the control cells (mean ±SD %c-Casp3 H2B-RFP+ cells control = 3.1 ±3.3 vs sh-SMAD3 = 4.0 ±1.9: Supp. Fig S2P; mean ±SD %pH3^+^ H2B-RFP+ cells control = 3.4 ±2.5 vs sh-SMAD3 = 3.4 ±2.1: Supp. Fig S2Q). Together, these observations prompted us to search for the precise cellular events regulated by TGF-β/SMAD3 signalling that might drive SNT formation.

### Confinement of NMPs and lineage restriction to NPCs are independent of Smad3 activity

Tail bud NMPs will generate NPCs in the SNT and trunk mesoderm. Co-expression of the T/Brachyury (T/Bra) and Sox2 transcription factors characterizes NMPs, while NPCs emerging from dual-fated NMPs downregulate T/Bra but maintain strong Sox2 expression (Gouti et al., 2014; Kondoh & Takemoto, 2012; Olivera-Martinez et al., 2012; Tsakiridis & Wilson, 2015; Wymeersch et al., 2016). Complete neural lineage restriction can be followed in stage HH15 chick embryos by analyzing T/Bra^+^ and Sox2^+^ expression at different axial levels, from the posterior elongating tail bud to the level of the last formed somite (Fig. 2A-E). At initial stages, T/Bra was expressed widely in the tail bud, even in the lateral mesoderm that was excluded from the quantifications here. Later in development, T/Bra was downregulated along the SN axis and finally, it became restricted to the notochord (Fig. 2A’-E’). Weak Sox2 expression initially appeared in the dorsal and central areas of the early tail bud (Fig. 2A’’) and this propagated ventrally (Fig. 2B’’) before eventually becoming confined to the NT (Fig. 2E’’). Hence, the differentiation of NPCs in the medullary cord apparently advances in a dorso-ventral direction.

**FIGURE 2:**
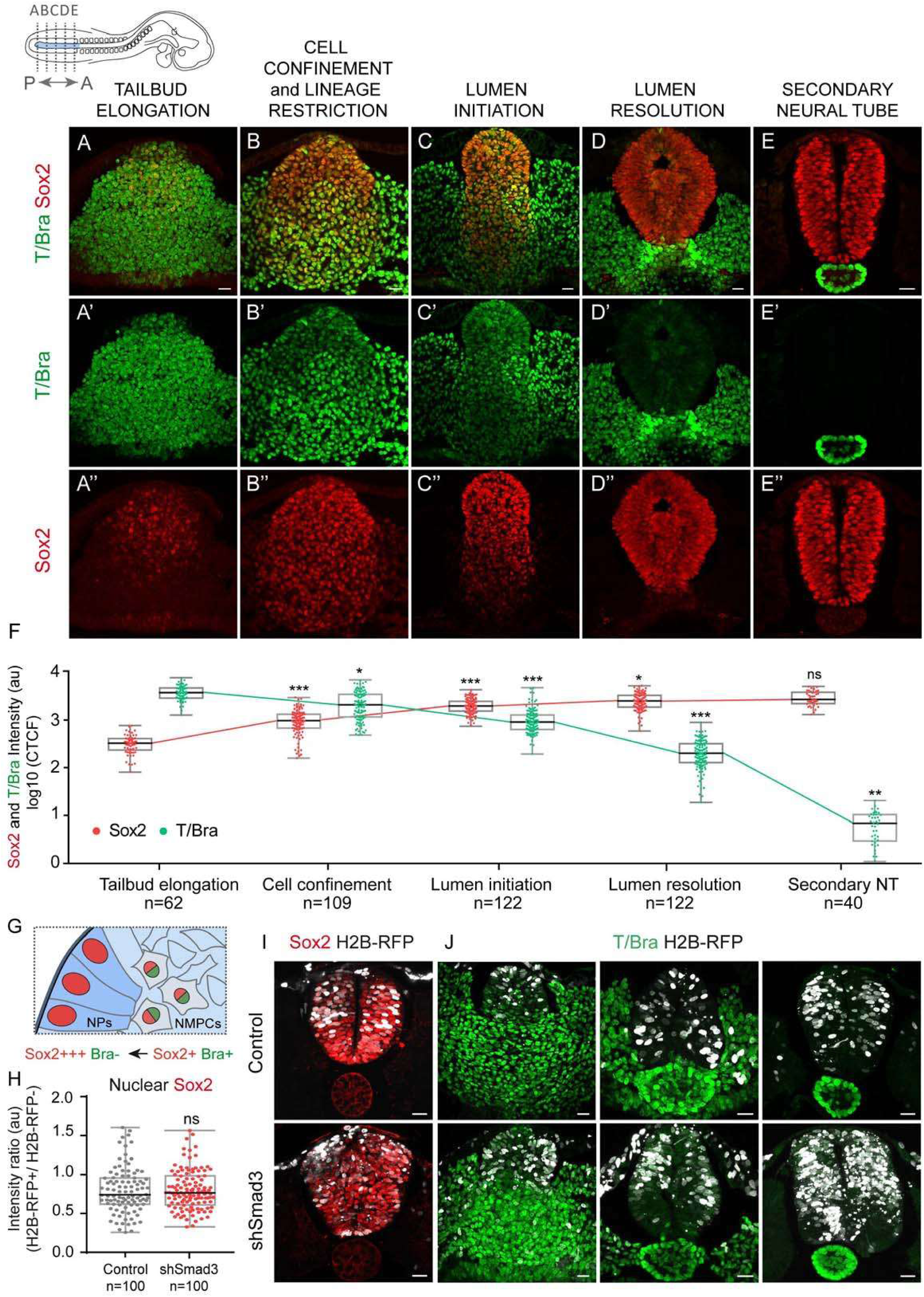
Confinement of NMPs and lineage restriction into NPCs are both independent of SMAD3 activity. **(A-E)** Selected images of transverse sections at the indicated cranio-caudal levels, stained for T/Bra (green) and Sox2 (red). Scale bars = 20 μm. **(F)** Nuclear fluorescence intensity of T/Bra and Sox2 during the indicated tissue remodelling events (horizontal bold lines show the median cell number from 10 embryos: n=62, 109, 122, 122, 40): *p<0.05, **p<0.01, ***p<0.001 Kruskal-Wallis test. **(G)** Scheme showing the developing SNT and lineage restriction of NMPs into SNT cells. Cells contacting the BM (purple) are the first to convert into PSNT cells, a transition characterised by T/Bra downregulation and Sox2 upregulation. **(H)** Plots ratio of Sox2 fluorescence intensity in control and sh-SMAD3 electroporated cells (bold horizontal lines show the median; n=100, 100 cells from 10 embryos/condition; *p>0*.*05* Mann-Whitney test). **(I)**Selected transverse sections of control and sh-SMAD3 electroporated embryos (white) showing the SNT stained for Sox2 (red). Scale bars = 20 μm. **(J)** Selected transverse sections of control and sh-SMAD3 electroporated embryos (white) showing the developing SNT stained for T/Bra (green). Scale bars = 20 μm.

Quantification of endogenous T/Bra and Sox2 within the region undergoing Secondary Neurulation not only revealed a strong reduction in T/Bra as this axis is established (median ±IQR log10 of T/Bra corrected total cell fluorescence (CTCF).axis elongation = 3.6 ±0.2; cell confinement = 3.3 ±0.5; lumen initiation = 3.0 ±0.3; lumen resolution = 2.3 ±0.4; SNT = 0.8 ±0.5: Fig. 2F), but also an increase in Sox2 (median ±IQR log10 of Sox2 CTCF axis elongation = 2.5 ±0.2; cell confinement = 3.0 ±0.3; lumen initiation = 3.3 ±0.2; lumen resolution = 3.4 ±0.2; SNT = 3.4 ±0.2: Fig. 2F), coincident with the progressive generation of NPCs. The expression of T/Bra and Sox2 shifted gradually as the SNT developed, suggesting the existence of a transitional state in the path from NMPs to NPCs. We also found a positive correlation between T/Bra expression and the distance from the basement membrane, and conversely, a negative correlation between Sox2 expression and its distance from the basement membrane (Supp. Fig. S3A,B). Moreover, during cell confinement, lumen initiation and lumen resolution, Sox2 was more strongly expressed by peripheral cells than central cells (median ±IQR log10 of Sox2 CTCF peripheral cells at cell confinement = 3.1 ±0.2; lumen initiation = 3.3 ±0.2; lumen resolution = 3.5 ±0.2 vs central cells at cell confinement = 2.9 ±0.4; lumen initiation = 3.2 ±0.2; lumen resolution = 3.3 ±0.2: Supp. Fig. S3C). Conversely, T/Bra expression was stronger in central cells than in peripheral cells (median ±IQR log10 of T/Bra CTCF peripheral cells at cell confinement = 3.1 ±0.3; lumen initiation = 2.9 ±0.3; lumen resolution = 2.2 ±0.4 vs central cells at cell confinement = 3.5 ±0.3; lumen initiation = 3.1 ±0.3; lumen resolution = 2.4 ±0.3: Supp. Fig. S3D). In summary, bipotential NMPs convert into NPCs by downregulating T/Bra and upregulating Sox2, which occurs in conjunction with cell confinement by the basement membrane (Fig. 2G).

To test the possible role of canonical TGF-β signalling in NMP lineage restriction, we further examined T/Bra and Sox2 expression 24 hpe of the sh-SMAD3 construct (Fig. 2H-J). T/Bra expression was progressively downregulated by NMPs electroporated with sh-SMAD3, although it was still expressed strongly by the surrounding lateral mesoderm cells, as well as by the axial notochord cells (Fig. 2J). The expression of Sox2 in NPCs electroporated with sh-SMAD3 was comparable to that in the surrounding non-electroporated cells (median ±IQR Sox2 intensity ratio control = 0.7 ±0.3 vs sh-SMAD3 = 0.8 ±0.4: Fig. 2H,I) and these Sox2^+^ cells became progressively restricted to the SNT (Supp. Fig. S3E). Moreover, by this stage (24 hpe of sh-SMAD3) a basal lamina clearly confined the SNT (Fig. 1L). All these observations indicate that SMAD3 activity does not affect cell confinement and that it is not required to achieve neural lineage restriction.

### Lumen initiation occurs at a one cell distance from the basement membrane and it is independent of SMAD3 activity

Concomitant with the neural lineage restriction of NMPs, these cells epithelialize as the SNT undergoes morphogenesis, a process that can be traced along the cranio-caudal axis of stage HH15 chick embryos. The first cells to acquire epithelial polarity are those in contact with the forming basement membrane, those located dorsally and at the periphery of the cord (Fig. 3A-D), and those undergoing epithelialisation similar to that of MDCK cells growing in 3D culture (Bryant et al., 2014; Martin-Belmonte et al., 2008) and in the early mouse embryo (Bedzhov & Zernicka-Goetz, 2014).

**FIGURE 3:**
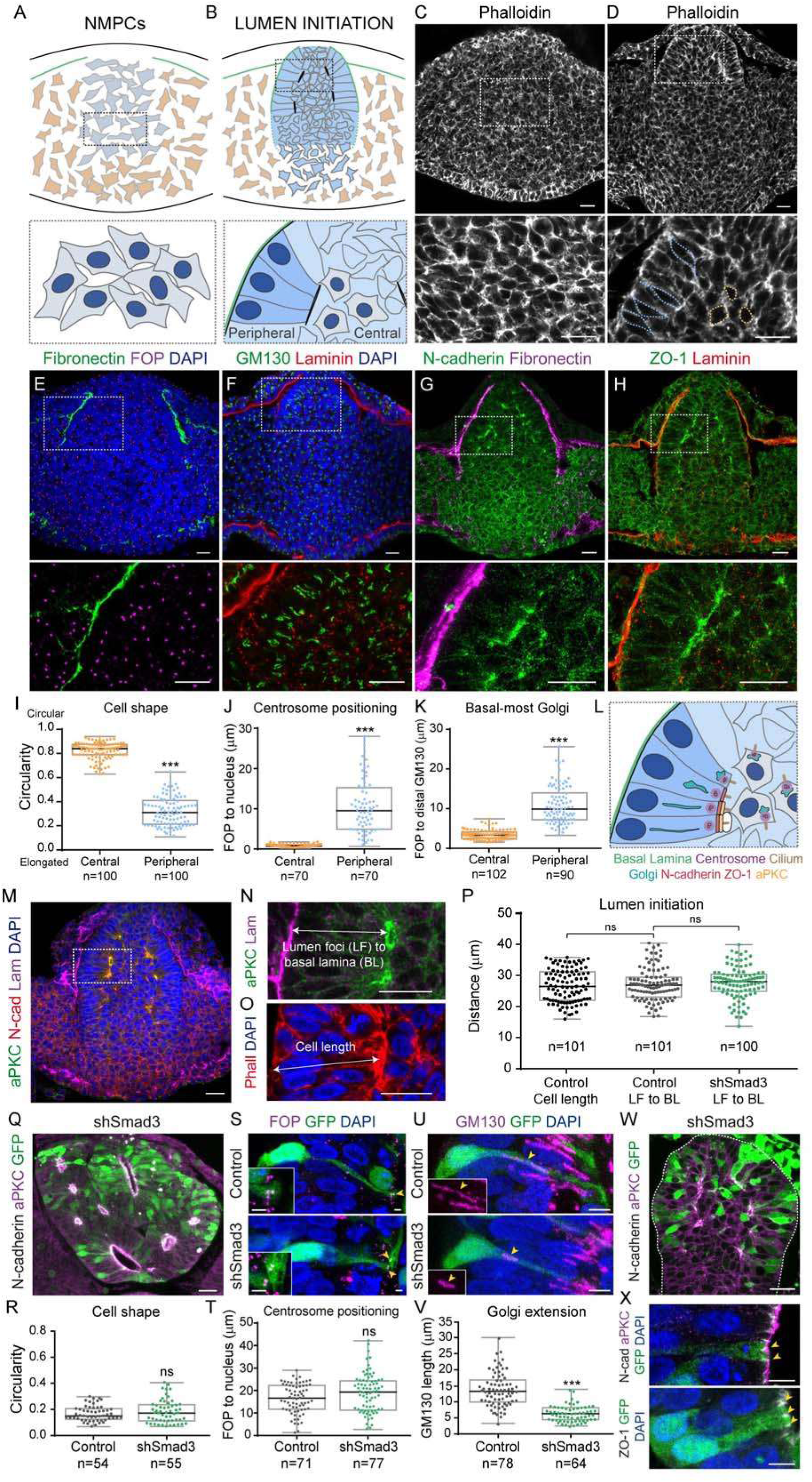
Lumen initiation occurs at an equivalent distance of one cell from the basal membrane and it is independent of SMAD3 activity. **(A**,**B)** Schemes representing the cellular processes occurring during MET of the NMPs (MET, mesenchymal-to-epithelial transition; NMPs, neuro-mesodermal progenitors). The developing basal membrane (BM) is in green, and in B, the central cells are shown in grey and the peripheral cells in light blue. **(C**,**D)** Selected transverse sections stained for actin (white). Higher magnifications of the boxed regions are shown in C’, D’ and the dotted lines in D’ define the central (orange) and peripheral (blue) cell shapes. Scale bars = 20 μm. **(E-H)** Selected transverse sections showing the markers indicated. A higher magnification of the boxed regions is shown in E’-H’. Scale bars = 20 μm. **(I)** Circularity plots of central and peripheral chord cells (bold horizontal lines show the median; n=100, 100 cells from 10 embryos; \**\*\*p<0*.*001* Mann-Whitney test). **(J)** Plots of the distance from FOP staining to the nucleus in central and peripheral chord cells (bold horizontal lines show the median; n=70, 70 cells from 10 embryos; \**\*\*p<0*.*001* Mann-Whitney test). **(K)** Plots of the distance from FOP to the basal-most part of the Golgi apparatus (distal GM130) in central and peripheral chord cells (bold horizontal lines show the median; n=102, 90 cells from 10 embryos; \**\*\*p<0*.*001* Mann-Whitney test). **(L)** Scheme of polarising PSNT cells. The centrosome is the first organelle to be apically localised, followed by the Golgi apparatus and N-cadherin/ZO-1, and finally aPKC. The BM (green) confines the cells undergoing MET. **(M)** Selected transverse sections at the stage of lumen initiation stained for aPKC (green), N-cadherin (red) and laminin (purple). A higher magnification of the boxed region appears in H’ to show the distance from lumen foci (LF) to the BM. Scale bars = 20 μm. **(N)** Selected transverse sections at the lumen initiation stage where cell length is visualised by actin (red) and nuclear (DAPI, blue) staining. Scale bar = 20 μm. **(O)** Selected transverse sections 24 hpe of sh-SMAD3 electroporation (green) at the lumen initiation stage showing the N-cadherin (white) and aPKC distribution (purple). Scale bar = 20 μm. **(P)** Plots of cell length and of the distance from the LF to BM in control and sh-SMAD3 embryos at the lumen initiation stage (bold horizontal lines show the median; n=101, 101, 100 cells from 10 embryos/condition; *p>0*.*05* Kruskal-Wallis test). **(Q)** Selected images of 24 hpe control and sh-SMAD3 electroporated cells (green) showing the actin cytoskeleton (white). Dotted pink lines delineate the cell shape, and the apical (a) and basal (b) surfaces are indicated for each cell. Scale bars = 5μm. **(R)** Plots cell shape/circularity in 24 hpe control and sh-SMAD3 electroporated embryos (bold horizontal lines show the median; n=54, 55 cells from 10 embryos/condition; *p>0*.*05* Mann-Whitney test). **(S)** Selected images of 2 4hpe control and sh-SMAD3 electroporated cells (green) showing the apically localised centrosomes (purple; yellow arrows). Higher magnifications are shown bottom left. Scale bars = 2.5μm. **(T)** Plots of the distance from the apical FOP to the nucleus in 24 hpe control and sh-SMAD3 electroporated embryos (bold horizontal lines show the median; n=71, 77 cells from 10 embryos/condition; *p>0*.*05* Mann-Whitney test). **(U)** Selected images of 24 hpe control and sh-SMAD3 electroporated cells (green) showing the *cis-*Golgi organisation (purple; yellow arrows). Scale bars = 5μm. **(V)** Plots of Golgi extension determined by GM130 staining of 24 hpe control and sh-SMAD3 electroporated embryos (bold horizontal lines show the median; n=71, 77 cells from 10 embryos/condition; \**\*\*p<0*.*001* Mann-Whitney test). **(W)** Selected images of sh-SMAD3 electroporated cells (green) stained for the indicated apical polarity proteins (yellow arrows). Scale bars = 5μm. **(X)** Selected transverse sections of embryos 24 hpe of sh-SMAD3 (green). N-cadherin (white) and aPKC (purple) line the small multiple lumens apically. Scale bar = 20 μm.

To characterize the subcellular events that accompany NMP epithelialisation, we analysed cell shape, the centrosome position, the Golgi distribution and the distribution of polarised proteins at the early stages of the MET (Fig. 3E-L, see also Supp. Fig. S4A-D). During MET in the SNT, the shape of the cells shifted from polygonal to elongated (median ±IQR circularity central cells = 0.8 ±0.1 vs peripheral cells = 0.3 ±0.2), concomitant with a basal translocation of the nucleus (Fig. 3E-I). The perinuclear centrosome relocated apically, as witnessed by its distance of the FOP^+^ labelled centrosomes (Yan et al., 2006) from the nucleus (median ±IQR distance central cells = 0.9 ±0.5 μm vs peripheral cells = 9.7 ±10.3 μm: Fig. 3E,J). The wide variation in the peripheral cell centrosome-to-nucleus distance is related to the onset of interkinetic nuclear migration, which separates or brings together the centrosome and the nucleus depending on the phase of the cell cycle. In addition, the Golgi elongated and became confined to the apical cellular process (median ±IQR distance central cells = 3.3 ±1.6 μm vs peripheral cells = 9.9 ±6.8 μm: Fig. 3F,K), as determined by labelling with the cis-Golgi matrix protein GM130 (Nakamura et al., 1995) and as observed for NPCs in the developing cerebral cortex (Taverna et al., 2016). Finally, cell epithelialization involves the reorganization of the apical membrane into discrete micro-domains where N-cadherin and the ZO-1/occludin complex occupy internal positions, whereas aPKC concentrates in the most apical domain (Aaku-Saraste et al., 1996; Afonso & Henrique, 2006; Chenn et al., 1998; Marthiens & ffrench-Constant, 2009). Indeed, apical proteins such as N-cadherin or ZO-1 progressively accumulated at the apical pole of the cell (Fig. 3G,H,L).

To establish the sequence of apical polarisation and the possible correlation with lumen initiation, we analysed the subcellular localisation of FOP, N-cadherin and aPKC during the initial stages of MET (Supp. Fig. S4E-M). While all the cells analysed already had an apically localised centrosome, several also presented apical N-cadherin and a few accumulated apical aPKC. Neither N-cadherin nor aPKC were distributed apically prior to apical centrosome positioning and similarly, apical aPKC was never detected prior to the apical accumulation of N-cadherin (mean ±sem % of peripheral cells with apical FOP = 73.7 ±5.1 vs apical FOP and N-cadherin = 26.3 ±5.1; apical FOP = 73.8 ±8.0 vs apical FOP and N-cadherin = 6.6 ±1.3 vs apical FOP, N-cadherin and aPKC = 19.6 ±8.9: Supp. Fig. S4E-I). Moreover, co-staining of FOP, N-cadherin and ZO-1 revealed that N-cadherin and ZO-1 co-localised apically at the same time, and again, never before FOP appeared apically (mean ±sem % of peripheral cells with apical FOP = 72.6 ±9.9 vs apical FOP, N-cadherin and ZO-1 = 27.4 ±9.9: Supp. Fig. S4E-I). Finally, in peripheral cells β1 Integrin was distributed basal before FOP accumulated apically, and no cells were found with only one of these two elements apico-basally polarised, which suggests that apical and basal polarity is organised concomitantly (mean ±sem % of peripheral cells with apical FOP and β1 integrin = 64.5 ±6.6 vs apical FOP, β1 integrin and N-cadherin = 35.5 ±6.6: Supp. Fig. S4H,J). In summary, we found that the subcellular events that lead to apico-basal polarisation of NPCs are sequential, whereby apical centrosome positioning and basal β1 integrin localisation is followed by N-cadherin/ZO-1 apical membrane accumulation, and finally, the apical membrane localization of aPKC (Fig. 3L).

As NMPs transformed into fully epithelialized NPCs, we observed multiple small lumens emerging at the interface between the peripheral epithelial and central mesenchymal cell populations (Fig. 3L,M). These small lumens always formed at a distance equivalent to one-cell from the basement membrane (median ±IQR cell length = 26.5 ± 9.2 μm vs LF to BM distance = 26.8 ±6.4 μm: Fig. 3M-P). We tested whether canonical TGF-β signalling might be required to trigger MET and to initiate lumen formation. At 24 hpe, sh-SMAD3 cells correctly restricted N-cadherin and aPKC to their apical pole, and multiple small lumens were established at one-cell distance from the BM (median ±IQR LF to BM distance control = 26.8 ±6.4 μm vs sh-SMAD3 = 28.0 ±5.5 μm: Fig. 3P). Hence, SMAD3 activity appears to be dispensable for the subcellular processes involved in triggering the MET and in the initial formation of lumen foci, which prompted us to search for defects in late stages of MET and SNT lumen resolution.

We tested whether canonical TGF-β signalling might be implicated in the final NPC epithelialization. To this end, we analysed cell shape, centrosome positioning, Golgi elongation and polar protein distribution in sh-SMAD3 electroporated cells in the SNT at the cranio-caudal levels where the morphogenesis of the neighbouring somites and underlying notochord had been completed. At 24 hpe, sh-SMAD3 cells of the SNT had properly elongated (median ±IQR circularity control = 0.1 ±0.1 vs sh-SMAD3 = 0.2 ±0.1: Fig. 3Q,R), and their centrosomes were situated apically, similar to control electroporated cells (median ±IQR FOP to nucleus distance control = 16.6 ±10.6 μm vs sh-SMAD3 = 19.3 ±13.0 μm: Fig. 3S,T). Although sh-SMAD3 electroporated cells failed to elongate their Golgi apparatus (median ±IQR GM130 length control = 13.3 ±6.9 μm vs sh-SMAD3 = 6.3 ±3.7 μm: Fig. 3U,V), these cells establish apical membrane micro-domains containing N-cadherin, ZO-1 and aPKC like control electroporated cells (Fig. 3W). Although, apico-basal polarity is correctly organized in NPCs, sh-SMAD3 electroporated embryos developed multiple lumens in the SNT, in which initial small lumens formed correctly but they failed to coalesce into a single central cavity (Fig. 3X). Together, these results indicated that SMAD3 activity was dispensable for the subcellular processes involved in cell epithelialization, yet it appeared to be required to establish a centrally positioned single lumen during SNT formation.

### SMAD3 activity is required for cell intercalation and the resolution of a single central lumen in the SNT

Formation of a single and continuous lumen in the SNT requires distinct cellular rearrangements and 3D tissue remodelling. 3D images of the lumen from ZO-1 *in toto* immunostained HH15 chick embryos were generated by Imaris reconstruction (Fig. 4A-F) to analyse their size and shape. Accordingly, the small caudal focal lumens were seen to first coalesce into three enlarged lumens (one dorsal-central and two ventral-lateral), which finally fused into a single central cavity in the more cranial domains (Fig. 4A-F, Sup Movie S1 and S2). We also found a population of cells that remained in between the lumens until they finally coalesced. Lumen resolution therefore required the clearance of this central cell population to generate a SNT composed of NPCs arranged around a single central cavity (Supp. Fig. S5A,I).

**FIGURE 4:**
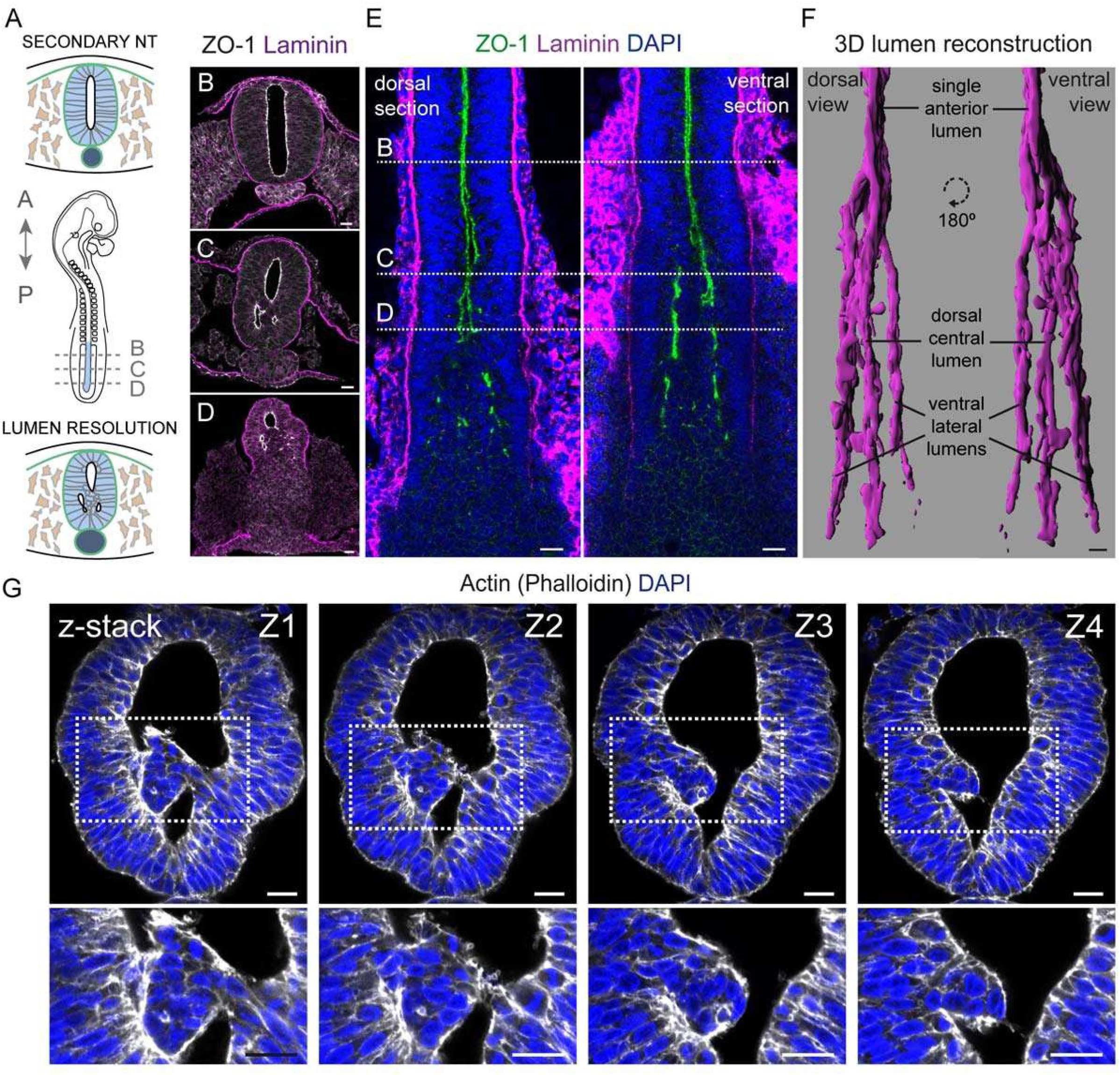
SMAD3 activity is required for cell intercalation and the resolution of a single central lumen in the SNT. **(A)** Drawing of a stage HH15 chick embryo and the cellular processes that occur during secondary lumen resolution to give rise to a SNT with a single central lumen. **(B-D)** Selected transverse sections at the indicated cranio-caudal levels in A, showing the localisation of the apical protein ZO-1 (white) and the basement membrane (BM, laminin, purple). The dotted yellow square encompasses the population of central cells. Scale bars = 20 μm. **(E)** Dorsal views at two different dorso-ventral levels of the SN region of a stage HH15 chick embryo stained for ZO-1 (green) and laminin (purple). DAPI (blue) stains cell nuclei and the dotted lines show the cranio-caudal levels in B-D. Scale bars = 20 μm. **(F)** Dorsal and ventral views of 3D reconstructions of the secondary lumen in a stage HH15 chick embryo. The single cranial lumen caudally splits into one dorsal-central lumen and two ventral-lateral lumens (n=5 embryos). Scale bar = 20 μm. **(G)** Consecutive sections of a transverse z-stack (Z1-Z4) at the lumen resolution phase with the actin (white) and nuclei (DAPI, blue) stained. Higher magnifications of the boxed regions are shown. Scale bars = 20 μm. **(H)** Selected transverse sections 24 hpe of sh-SMAD3 (green) at the lumen resolution stage, stained for ZO-1 (white). A higher magnification of the boxed region is shown in E’. Scale bars = 20 μm. **(I)** Selected transverse sections 24 hpe of sh-SMAD3 (green) with actin (white) and the cell nuclei (DAPI, blue) stained. A higher magnification of the boxed region is shown in F’. Scale bars = 20 μm.

The analysis of these central cells revealed that even though they had already lost T/Bra expression (Supp. Fig. S5A,B) and were therefore referred to as central NPCs, they retained certain mesenchymal characteristics. Central NPCs remained round at a point when peripheral NPCs at that stage of lumen resolution had already lost their circularity and were heading towards full elongation in the formed SNT (median ±IQR circularity central NPCs (LR) = 0.8 ±0.1, peripheral NPCs (LR) = 0.2 ±0.1, NPCs (SNT) = 0.1 ±0.03: Supp. Fig. S5C,F). The centrosome of these central NPCs also remained close to their nucleus (median ±IQR FOP to nucleus distance central NPCs (LR) = 1.1 ±0.6 μm, peripheral NPCs (LR) = 14.3 ±11.0 μm, NPCs (SNT) = 27.7 ±21.2 μm: Supp. Fig. S5D,E,G), and they retained a pericentrosomal Golgi apparatus (median ±IQR FOP to distal GM130 distance central NPCs = 4.8 ±2.8 μm, peripheral NPCs = 16.8 ±10.3 μm, NPCs (SNT) = 22.2 ±20.3 μm: Supp. Fig. S5D,E,H) and their apical polarity was disorganised (Supp. Fig. S5E).

Resolution into a single continuous lumen in the SNT requires the clearance of the central mesenchymal NPCs. To test whether cavitation is implicated in this lumen resolution, cleaved caspase-3 and TUNEL staining was performed on stage HH15 chick embryo sections at various cranio-caudal levels. While a variable numbers of apoptotic cells were detected in the dorsal NT and dorsal non-neural ectoderm, such cells were almost virtually absent from the forming SNT (mean ±sem %c-Caspase3 NPCs = 0.6 ± 0.1), or even from the inner cell mass surrounded by forming lumens (mean ±sem %c-Caspase3 central NPCs = 0.3 ±0.2), indicating that cavitation does not contribute to lumen resolution in the SNT (Supp. Fig. S5M-O). However, the population of centrally located NPCs appeared to intercalate among the epithelialized NPCs of the developing SNT, as observed in transverse sections in which actin was stained with Phalloidin (Fig. 4G). Interestingly, this process of lumen resolution by cell intercalation appeared to require SMAD3 activity, since at 24 hpe, sh-SMAD3 electroporation led to the formation of centrally located NPCs at lumen resolution stages that did not undergo apoptosis and die (Supp. Fig. S2M,N,P).

To directly test this hypothesis, we electroporated stage HH9 chick embryos to analyse the process *in vivo*, which were later cultured and imaged under an upright wide-field microscope. This system allows chick embryos to elongate and to develop at the approximate same rate as they do *in ovo* (Benazeraf et al., 2010; Gonzalez-Gobartt et al., 2020; Rupp et al., 2003). It also permits fluorescently labelled electroporated cells in the elongating SNT to be tracked over time. These cells are highly motile, with a high rate of cell division, and they undergo important cell mixing. Analysis of membrane-GFP electroporated cells revealed round mesenchymal cells situated centrally in the elongating SNT (green arrows in Fig. 5Ai and first time-point in D-F). We consistently observed central cell intercalation into the lateral walls of the neuroepithelium, perceived as a lateral cell movement away from the centre of the tissue, and this was accompanied by an elongation of the cells (Fig. 5A-C; Supp. Movies S3). Notably, intercalating cells were highly protrusive (Fig. 5A-E) and their intercalation always occurred just after cell division (dotted circles in Fig. 5B-F). Indeed, the proportion of mitotic cells in fixed sections of stage HH15 embryos was significantly higher in the area where the single lumen is resolving than in the formed SNT or the caudal tail bud (mean ±SD % pH3 cells SNT = 4.4 ±1.0 μm vs LR = 7.0 ±2.5 μm vs Tail bud = 3.3 ±1.1 μm: Fig. 5G).

**FIGURE 5:**
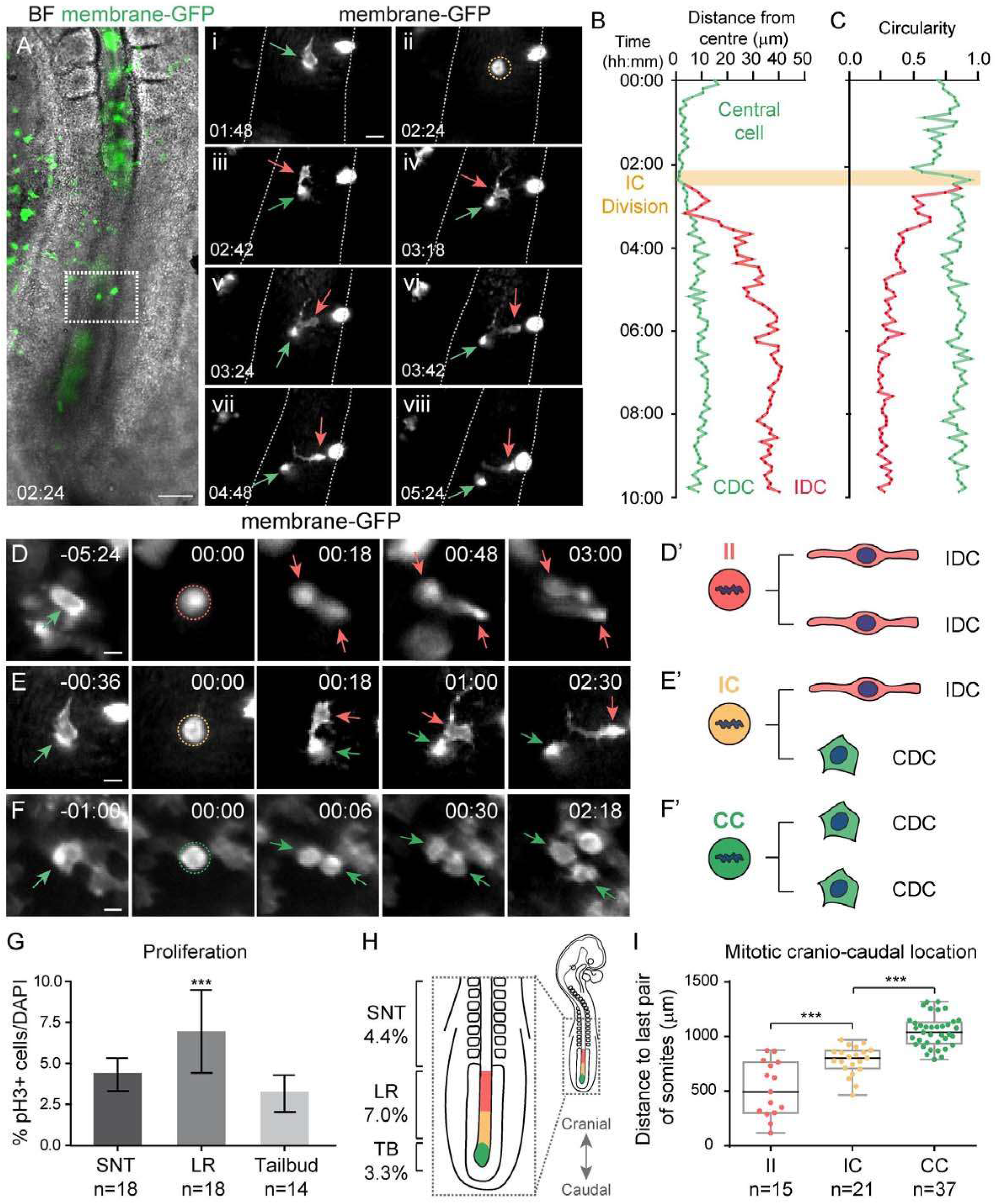
Three modes of division co-exist in the developing SNT but at different cranio-caudal levels. **(A)** Selected frames of membrane-GFP electroporated embryos time-lapse movies showing a representative intercalating (IC) mitosis. A central mesenchymal cell (i, green arrow) divides (ii, orange dotted circle) and generates one intercalating daughter cell (IDCs: iii-viii, red arrow) and one central daughter cell (CDC: iii-viii, green arrow). Scale bars = 100 μm, 20 μm. **(B)** Distance from the centre of the cells in D during the time-lapse movie (10 h). One daughter cell remains in the centre (green) while the other moves away and intercalates into the lateral wall of the SNT (red). **(C)** Circularity of the cells in D during the time-lapse movie (10 h). One daughter cell maintains the high circularity of the mother cell (green) while the other elongates (red). **(D-F)** Selected frames of time-lapse movies of membrane-GFP electroporated embryos showing a representative mitosis for each of the three modes of cell division. The parental central mesenchymal cell (green arrow) generates two IDCs (dark and light red arrows) in A; one CDC (green arrow) and one IDC (red arrow) in B; and two CDCs (dark and light green arrows) in C. Scale bars = 10 μm. **(D’-F’**) Schemes of the three modes of division that coexist during SNT formation. A cell undergoing mitosis can generate either two IDCs (D’, II mode of division, red), one IDC and CDC (E’, IC mode of division, yellow) or two CDCs (F’, CC mode of division, green). **(G)** Plots of the mitotic pH3^+^ cells once SNT is formed, in the lumen resolution stage (LR) and in the early elongating tailbud (TB, mean ±SD n = 18, 18, 14 sections from 10 embryos: **p<0.01,***p<0.001 one-way ANOVA). **(H)** Scheme of a stage HH15 chick embryo showing a higher magnification of its caudal region. The mean mitotic indexes in D are indicated for each region (left). The approximate cranio-caudal location of each mode of division is also shown (II, red; IC, yellow; CC, green). **(I)** Plots of the cranio-caudal distribution of each of the three modes of division represented as the distance of each division to the last formed pair of somites (bold horizontal lines show the median; n=15, 21, 37 divisions from 13 time-lapse movies; ***p<0.001 one-way ANOVA).

We quantified the distance of central mitotic membrane-GFP^+^ cells from the centre, their circularity and cranio-caudal location (relative to the last pair of somites formed) over time, and that of their daughter cells. The outcome of central cell division varied along the cranio-caudal axis (Fig. 5D-F). At the anterior SNT (median ±IQR distance to the last pair of somites = 494.8 ±464.0 μm), central cells divided symmetrically before both daughter cells elongated and migrated laterally to intercalate among the epithelialized NPCs (Fig. 5D,H,I; Supp. Movie S4). We referred to this mode of division as II, as it produced two intercalating daughter cells (IDCs: Fig. 5D’). In the intermediate regions (median ±IQR distance to the last pair of somites = 803.2 ±163.2 μm), central cells divided asymmetrically, such that one daughter cell remained mesenchymal and centrally located, while the other cell elongated and migrated laterally to intercalate among epithelialized NPCs forming the SNT (Fig. 5E,H,I; Supp. Movie S3). We termed this mode of division as IC, as it gave rise to one IDC and a central daughter cell (CDC: Fig. 5E’). At the caudal tail bud (median ±IQR distance to the last pair of somites = 1040.0 ±193.1 μm) central mesenchymal cells divided symmetrically, such that both daughter cells remain round and centrally located (Fig. 5F,H,I; Supp. Movie S5). We termed this mode of division as CC, as it generates two CDCs (Fig. 5F’).

To test whether canonical TGF-β signalling influenced cell intercalation and lumen resolution during SNT formation, stage HH9 chick embryos were electroporated with a sh-SMAD3 or control Sox2p:GFP vector (Saade et al., 2013; Uchikawa et al., 2003) to track the fluorescently labelled cells over time (Fig. 6A-D). Each of the three modes of division were evident at the corresponding cranio-caudal levels in Sox2p:GFP control embryos (median ±IQR distance to the last pair of somites control II = 732 ±378.8 μm, IC = 843 ±123 μm, CC = 1008 ±163 μm: Fig. 6E) and daughter cells intercalated normally (Supp. Fig. S6J,K; Supp. Movie S6). Although sh-SMAD3 electroporated central cells divided in a similar ratio as the control electroporated cells (Supp. Fig. S2M,O,Q), most divisions generated two cells that remained centrally located in sh-SMAD3 electroporated embryos. These daughter cells failed to intercalate into the lateral epithelialized SNT (Fig. 6F-I; Supp. Movies S7, S8), regardless of their mitotic cranio-caudal location (median ±IQR distance to the last pair of somites sh-SMAD3 IC = 718.8 ±503.5 μm, CC = 831.9 ±445.8 μm). Faulty cell intercalation in sh-SMAD3 electroporated embryos resulted in the abnormal persistence of central cells at cranial levels (Fig. 6C,D), that eventually elongated at these central positions (last time points in Fig. 6G-I). These alterations disturbed the resolution of the developing lumen and ultimately led to NTDs involving a multi-lumen phenotype.

**FIGURE 6:**
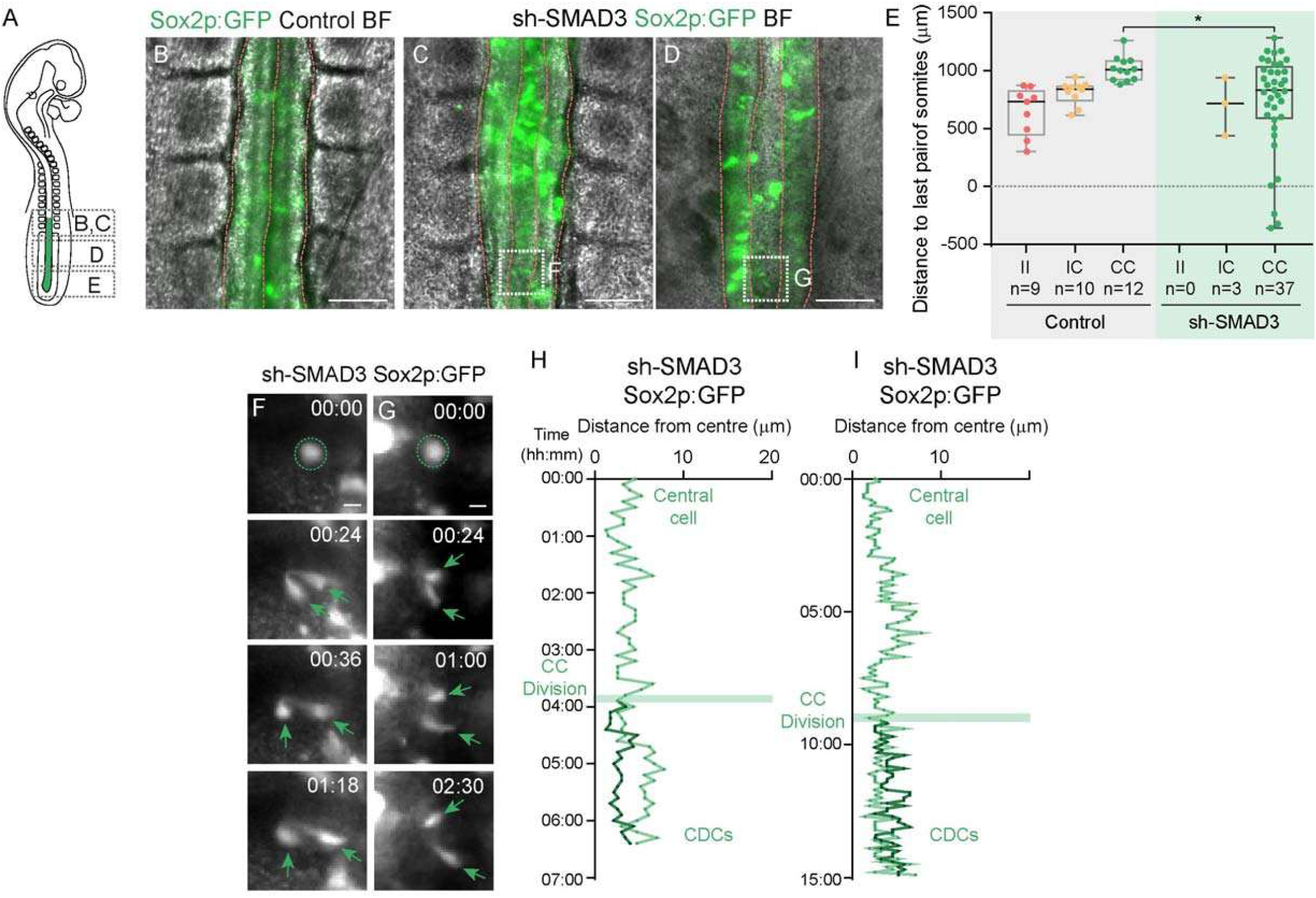
sh-SMAD3 electroporated cells fail to intercalate into the lateral walls of the forming SNT. **(A)** Drawing of an electroporated HH15 chick embryo showing the cranio-caudal levels of B-E. **(B)** Selected frame of Sox2p:GFP electroporated control embryos time-lapse movies at the cranio-caudal level indicated in A. Red dotted lines define the apico-basal surfaces of the SNT. Scale bar = 100 μm. **(C-E)** Selected frames of time-lapse movies of sh-SMAD3 Sox2p:GFP electroporated control embryos at the cranio-caudal levels indicated in A. Frames are the last time points of the mitotic sequences in G,H,I, as indicated by the boxed regions. Red dotted lines define the apico-basal surfaces of the SNT. Note the abnormal accumulation of cells in the centre of the SNT in C and D. Scale bars = 100 μm. **(G-H)** Selected frames of sh-SMAD3 Sox2p:GFP movies showing two different cell divisions at the cranio-caudal positions indicated in C,D. The dotted circles indicate the mitosis (CC divisions, green), the green arrows point to central daughter cells (CDCs) and the red arrow to an intercalating daughter cell (IDC). Scale bars = 10 μm. **(F)** Plots of the cranio-caudal location for each of the three modes of division as the distance from each division to the last formed pair of somites in both Sox2p:GFP control and sh-SMAD3 electroporated embryos. Almost all sh-SMAD3 divisions generate two CDCs (CC divisions) regardless of the cranio-caudal position (bold horizontal lines show the median; n=9,10,12 divisions from 5 Sox2p:GFP control and n=0,3,37 divisions from 6 sh-SMAD3 time-lapse movies): *p<0.05 Kruskal-Wallis test. **(J)** Selected frame of Sox2p:GFP control time-lapse movies. The boxed region refers to the last time point (v) of the sequence in B. Scale bar = 100 μm. **(K)** Selected frames of HH15 Sox2p:GFP time-lapse movies showing a representative IC mitosis. A mesenchymal cell (i, green arrow) divides (ii, orange dotted circle) and generates a CDC (iii-v, green arrow) and an IDC that intercalates into the lateral wall of the SNT (iii-v, red arrow). Scale bars = 10 μm. **(L)** Plots of the cell distance from the centre of the cells in B during the time-lapse movie. One daughter cell remains in the centre (green) while the other moves away and intercalates into the lateral wall of the SNT (red).

## DISCUSSION

In this study, we provide an exhaustive description of the morphogenetic events through which the SNT is generated in the chick embryo, information that will be crucial to unravel the causes of caudal NTDs, among the most common birth defects in humans. Importantly, we found that the final resolution of the central lumen involves cell intercalation into the lateral walls of the neuroepithelium, leading to a revision of the pre-existing model of secondary neurulation in the chick embryo (Schoenwolf & Delongo, 1980). We also show that defective TGF-β/SMAD3 activity during the formation of the SNT leads to caudal NTDs characterised by the presence of multiple small lumens. This phenotype is not due to changes in cell viability, cell identity, lumen initiation or apico-basal polarity disruption but rather, it arises from a failure in central cell intercalation during lumen resolution.

We studied the process of secondary neurulation starting from the stage when mesenchymal NMPs drive caudal body axis elongation to the complete formation of the SNT, in which epithelial NPCs surround a single central lumen. Based on our findings, we divided the morphogenesis of the SNT into three fundamental steps: (i) confinement of NMPs and neural lineage restriction; (ii) MET and *de novo* formation of multiple lumens; (iii) lumen resolution into a single central cavity. In our model SNT formation involves two related cell events, a change in cell identity from NMPs to NPCs and subsequently, the polarization of mesenchymal (front-rear) into epithelial (apico-basal) cells. Both transformations are associated with the growth of the basement membrane, which suggests an important role for basement membrane/integrin signalling in this process. Concomitant to these cellular changes, the SNT lumen forms *de novo* in between cells, with small lumens opening up at the equivalent of a one-cell distance from the basement membrane. The formation of these lumens is associated with the isolation of a central mesenchymal cell population. The accepted model for SN does not explain how this population of central cells is cleared from the lumen, although the possibility of central cell intercalation into the lateral walls of the neuroepithelium has been contemplated (Schoenwolf & Delongo, 1980). However, although this hypothesis was proposed decades ago, it has yet to be tested. By performing *in vivo* time-lapse imaging in electroporated chick embryos, we were able to follow central NPCs over time, showing that central cells indeed intercalate into the lateral walls of the neuroepithelium as the lumens initially formed coalesce. The result of this process is that a hollow SNT is formed, composed of epithelial NPCs encompassing a single central lumen. Together, we bring together data on cell identity, cell polarity and cell dynamics to extend and complete the pre-existing model of normal SNT development, establishing the basis to understand the pathology of caudal NTDs.

The molecular signals driving SNT formation remain largely unknown. WNT and FGF signalling play important roles in the maintenance and expansion of NMPs (Garriock et al., 2015; Takemoto et al., 2006; Wymeersch et al., 2016; Yamaguchi et al., 1999), and in the induction of the mesodermal or neural lineage of NMP derivatives (Diez del Corral et al., 2002; Gouti et al., 2017; Martin & Kimelman, 2012; Nowotschin et al., 2012; Yoshikawa et al., 1997). Here we show that both TGF-β and BMP signalling pathways are active during SNT formation, supporting the results obtained from our analysis of gene expression. While normal SNTs form even when BMP SMADs are inhibited, depletion of TGF-β SMAD3 but not SMAD2 generates a NTD characterised by a multi-lumen phenotype. We cannot completely rule out a role for BMP in this process, as the negative results obtained with BMP sh-SMAD electroporation could reflect a requirement for the inhibition of the BMP pathway for the SNT to form. The different outcomes following the inhibition of two TGF-β SMADs might reside in the fact that SMAD3 and SMAD2 can either co-operate or antagonize each other to regulate their transcriptional targets (Miguez et al., 2013). Phosphorylated R-SMAD proteins form heterotrimeric complexes with SMAD4 that enter the nucleus, where they recruit various co-factors and bind to DNA in order to regulate target gene expression (A Moustakas et al., 2001; Aristidis Moustakas & Heldin, 2002). These complexes can contain either SMAD2 or SMAD3 homodimers, or SMAD2/SMAD3-heterodimers (Shi & Massagué, 2003). As SMAD2 and SMAD3 recruit different co-factors and target different regulatory sequences (Brown et al., 2007), the balance between the heterotrimeric complexes formed upon ligand stimulation dictates the set of genes that will ultimately be activated. Our results therefore indicate that the activation of SMAD3 targets but not that of SMAD2 targets, is crucial for SNT morphogenesis.

TGF-β signalling is known to participate in the disruption of epithelial organisation and EMT (Yang & Weinberg, 2008). However, we show here that SMAD3 depletion does not affect the capacity of cells to polarise apico-basally and initiate the formation of multiple lumens as part of the development of the SNT. Indeed, SMAD3 deficient cells normally elongate and correctly localise their centrosomes to the apical pole. The Golgi apparatus is the only organelle that we found to be altered in SMAD3 deficient NPCs, as it fails to extend as in control cells. Although the Golgi apparatus is abnormally short, it remains confined to the apical process of NPCs, between the nucleus and the apical membrane. It is important to note that we only studied the cis-Golgi through GM-130 staining, leaving the trans-Golgi unexplored. Nevertheless, the Golgi apparatus of NPCs in the developing neocortex is oriented with their cis-to-trans axis perpendicular to the apico-basal axis of the cell (Taverna et al., 2016). If this orientation were conserved in spinal cord NPCs, then the extension of the Golgi will not affect apico-basal trafficking to a large extent but rather, it will be influenced by other Golgi cisternae functionalities. Indeed, junctional and apical markers like N-cadherin, ZO-1 and aPKC adopt a normal sub-cellular distribution and completely line the multiple luminal surfaces, suggesting that apical intracellular trafficking occurs normally. The fact that depletion of TGF-β activity results in a multi-lumen phenotype with correct apico-basal polarity, as opposed to the absence of polarisation and the complete inexistence of a lumen, points to a role of this pathway in the last steps of SNT lumen formation.

We found that the TGF-β pathway is active and SMAD3 is expressed strongly by central cells at the stage of lumen resolution. We hypothesize that TGF-β signalling, through SMAD3, could replace the lost T/Bra activity in central NPCs and ensure that they retain their mesenchymal features. The mesenchymal capacities retained confer high cell motility and invasive properties to central cells, permitting their intercalation into the lateral walls of the developing neuroepithelium. Indeed, TGF-β activity controls actin polymerisation and actomyosin contractility during lumen expansion in the Ciona intestinallis notochord (Denker et al., 2015). Both actin polymerisation and actomyosin contractility are essential for mesenchymal cell migration (Chi et al., 2014; Ridley et al., 2003). Moreover, TGF-β activity also controls stress fibre formation, the reorganization of cytoskeletal structures and the mesenchymal characteristics of cells through the activation of RhoA-dependent signalling (Bhowmick et al., 2001; Edlund et al., 2002; Shen et al., 2001). However, the precise mechanisms by which TGF-β/SMAD3 regulates the motility of central cells and promotes cell intercalation remain to be determined.

Together, we propose here a model in which central cells lose their motility and their protrusive behaviour in the absence of SMAD3 activity, such that they become unable to clear from central areas and intercalate. These data indicate that SMAD3 deficiency prevents central lumen resolution, ultimately generating NTDs.

## ACKNOWLEDGMENTS

The authors are indebted to Dr Elena Rebollo for her invaluable technical assistance at the AFMU Facility (IBMB); We thank Leica Microsystems for supporting and collaborating with the AFMU Facility (IBMB). We are grateful to researchers that kindly provided DNAs and antibodies, as indicated in the reporting summary. The work in EM’s laboratory was supported by grants BFU2016-77498-P, BFU2016-77498-P and La Maratò de TV3 foundation 201833-10.

## AUTHOR CONTRIBUTIONS

**EGG** conceived and performed most of the experiments, analysed the data, discussed the results and revised the manuscript.

**JBA** contributed to the experiments, the image acquisition, image analysis and quantification, and the statistics, and they revised the manuscript.

**SU** provided technical support for most experiments and performed in situ hybridization screen.

**GA and BB** conceived the in vivo imaging experiments, discussed the results and revised the manuscript.

**EM** conceived the experiments, analysed the data, discussed the results and drafted the manuscript.

## FIGURE LEGENDS

**SUPPLEMENTARY FIGURE S1:**
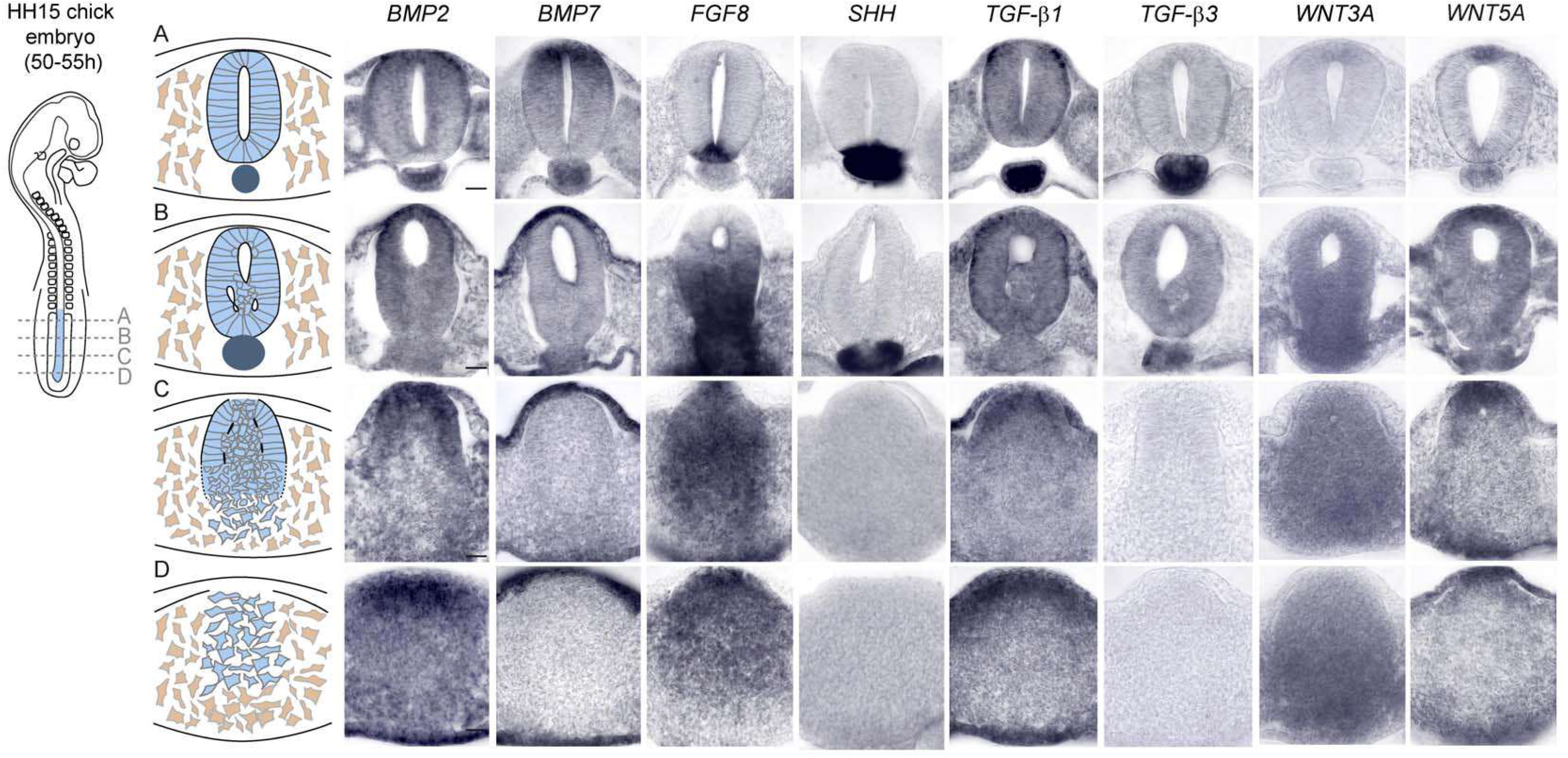
Analysis of the mRNA expression of potential morphogenetic signals in the forming SNT. Transverse sections of embryos subjected to whole mount in situ hybridization to assess the expression of *mRNAs* for several secreted proteins. Selected transverse sections at cranio-caudal levels A-D hybridized with probes for the *mRNAs* indicated. Scale bars = 20 μm.

**SUPPLEMENTARY FIGURE S2:**
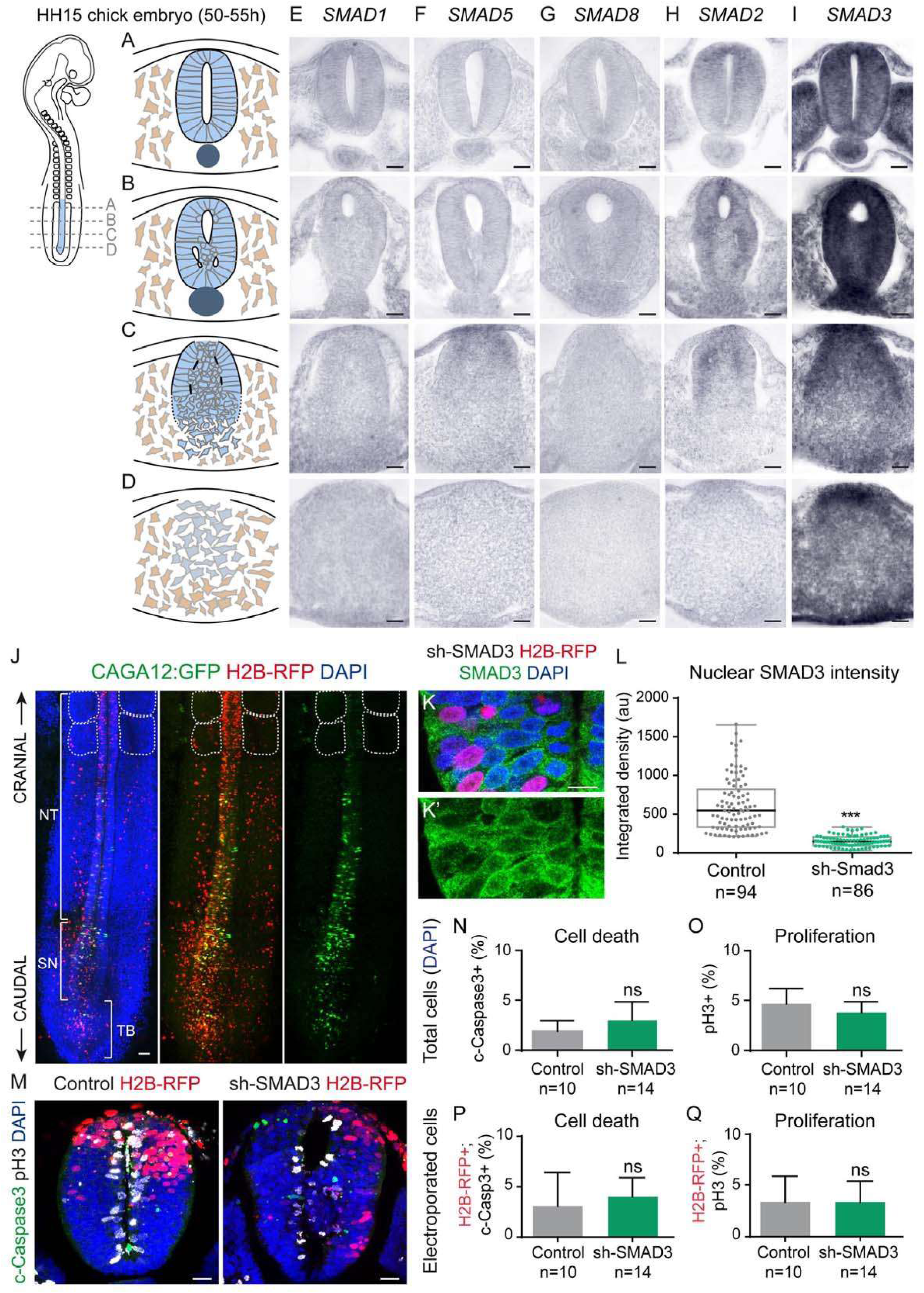
Analysis of SMAD expression and activity in the forming SNT. Transverse sections of embryos subjected to whole mount in situ hybridization tto assess the expression of SMAD *mRNAs*. Selected transverse sections at cranio-caudal levels A-D hybridized with probes for the *mRNAs* indicated. Scale bars = 20 μm. **(E**,**F**,**G)** BMP signalling SMADs. **(H**,**I)** TGF-β signalling SMADs. **(J)** Dorsal view of CAGA12:GFP (green) and control H2B-RFP (red) embryos 24 hpe. DAPI (blue) stains the nuclei and the dotted lines define the somites: NT, neural tube; SN, secondary neurulation region; TB, tailbud. Scale bar = 40 μm **(K)** Selected transverse sections 24 hpe of sh-SMAD3 and control H2B-RFP (red). DAPI (blue) stains the cell nuclei and anti-SMAD3 (green) stains endogenous SMAD3 (protein quantification plotted in L). Scale bars = 40 μm. **(L)** Nuclear fluorescence intensity in control and sh-SMAD3 electroporated cells (bold horizontal lines show the median; n=94, 86 cells from 10 embryos/condition): ***p<0.001 Mann-Whitney test. **(M)** Selected transverse sections 24 hpe of control or sh-SMAD3, together with control H2B-RFP (red). DAPI (blue) stains the cell nuclei, pH3 (white) the dividing cells and c-Caspase3 (green) the apoptotic cells. Scale bars = 20 μm. **(N)** Plots the proportion of c-Caspase3^+^ cells (relative to the total cell number -DAPI) in control and sh-SMAD3 electroporated embryos (mean ±SD, n=10-14 embryos/condition): p>0.05 Mann-Whitney test. **(O)** Proportion of mitotic pH3^+^ cells relative to the total cell number -DAPI) in control and sh-SMAD3 electroporated embryos (mean ±SD, n=10-14 embryos/condition): p>0.05 Mann-Whitney test. **(P)** Proportion of c-Caspase3^+^/H2B-RFP^+^ cells (relative to the total H2B-RFP^+^ cells) in control and sh-SMAD3 electroporated embryos (mean ±SD, n=10-14 embryos/condition): p>0.05 Mann-Whitney test. **(Q)** Proportion of mitotic pH3^+^/H2B-RFP^+^ cells (relative to the total H2B-RFP^+^ cells) in control and sh-SMAD3 electroporated embryos (mean ±SD, n=10-14 embryos/condition): p>0.05 Mann-Whitney test.

**SUPPLEMENTARY FIGURE S3:**
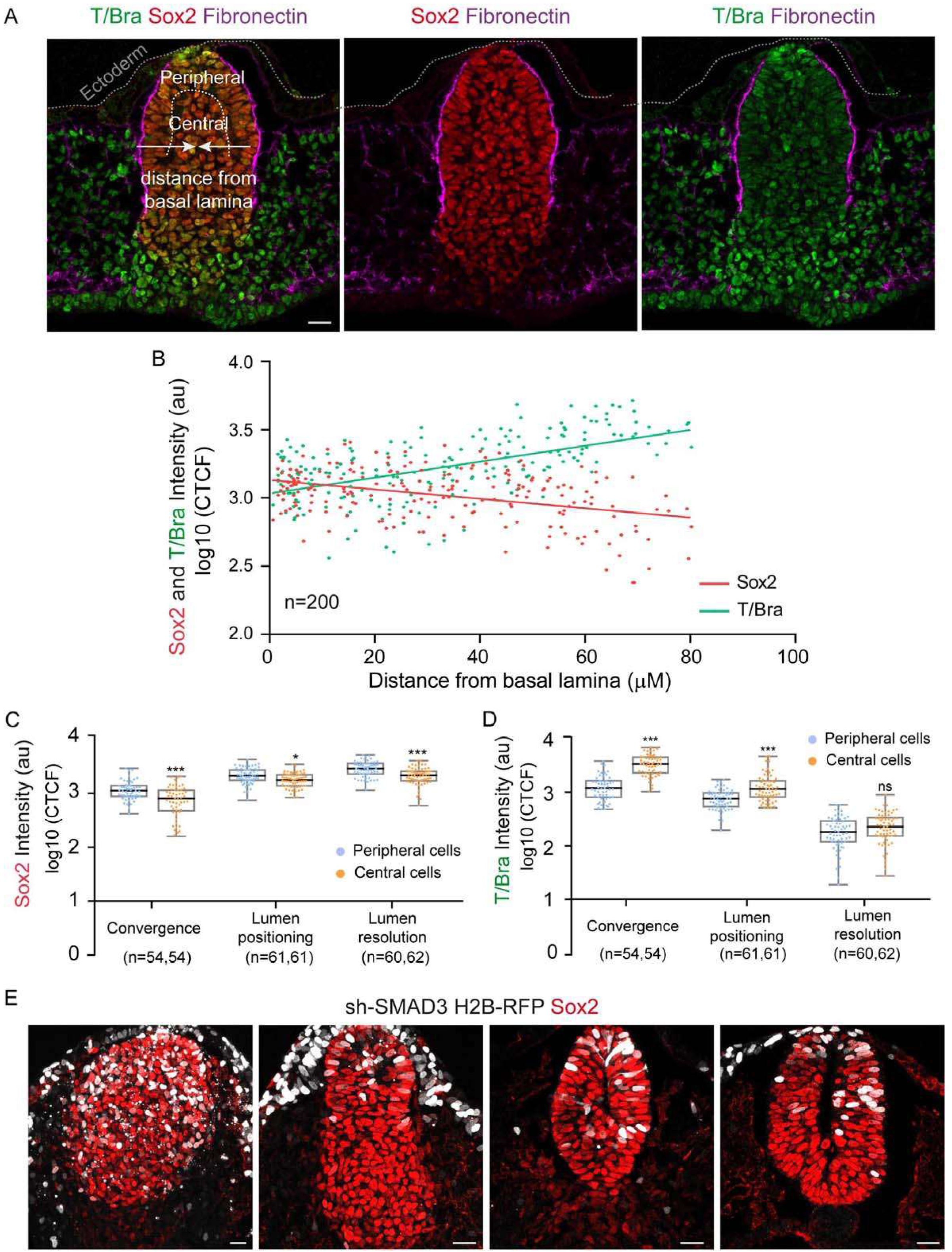
Lineage restriction of NMPs in association with confinement by the basal membrane. **(A)** Selected transverse sections stained for T/Bra (green), Sox2 (red) and Fibronectin (purple). A dotted white line separates the peripheral and central cell populations. The distance of the cell nucleus from the basement membrane (BM) is calculated. Scale bar = 20 μm. **(B)** T/Bra and Sox2 nuclear fluorescence intensity at the distance from the BM indicated. **(C)** Nuclear T/Bra fluorescence intensity in central and peripheral NPCs in association with the tissue remodelling events indicated (bold horizontal lines show the median; n=54,54; 61, 61; 60,62 cells from 10 embryos): ***p<0.001 Kruskal-Wallis test. **(D)** Nuclear Sox2 fluorescence intensity in central and peripheral NPCs in association with the tissue remodelling events indicated (bold horizontal lines show the median; n=54,54; 61, 61; 60,62 cells from 10 embryos): *p<0.05, ***p<0.001 Kruskal-Wallis test. **(E)** Selected transverse sections of control and sh-SMAD3 electroporated embryos (white) showing the developing SNT stained for Sox2 (red). Scale bars = 20 μm.

**SUPPLEMENTARY FIGURE S4:**
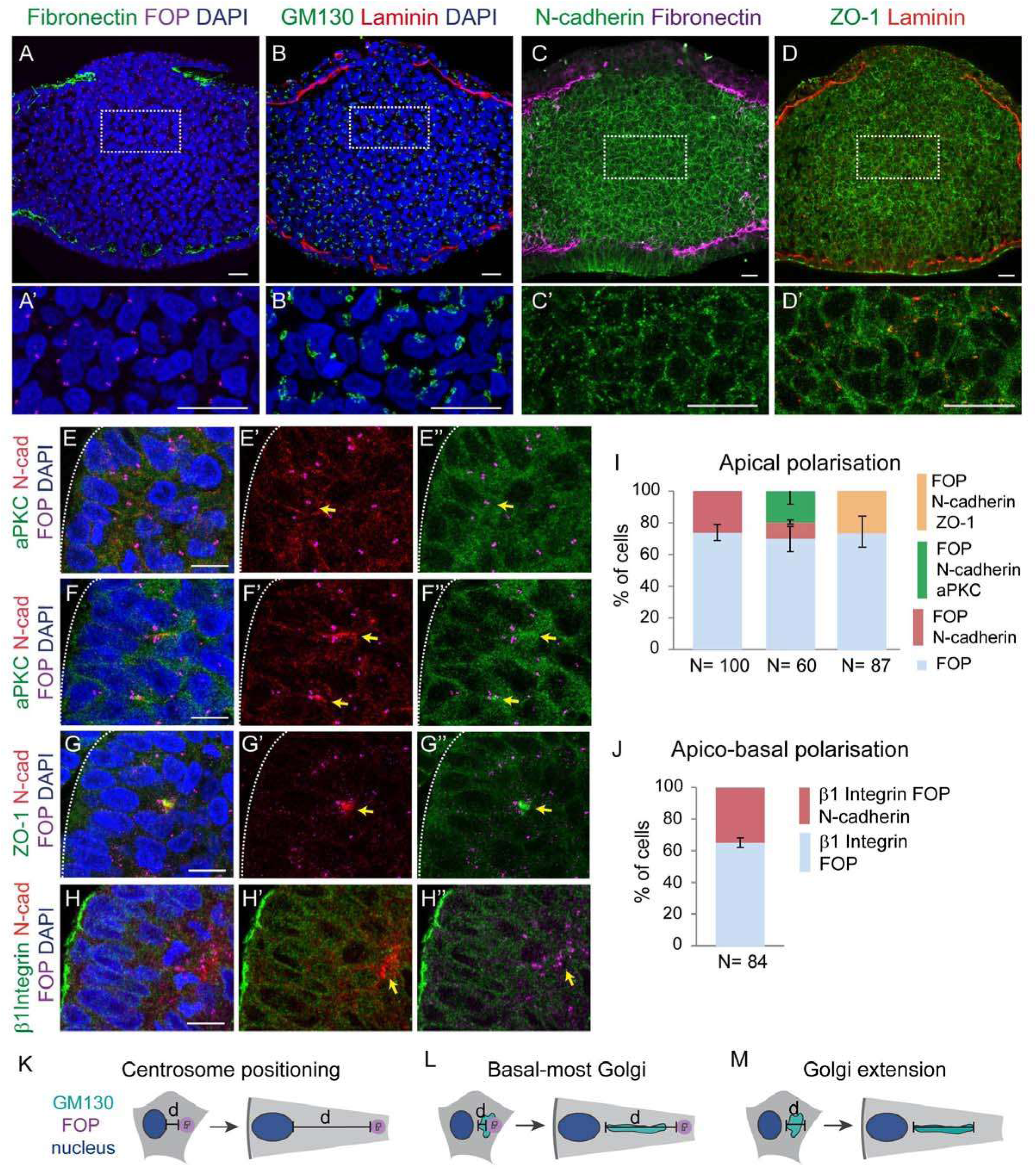
The Mesenchymal-to-epithelial transition occurs in association with the basement membrane. **(A)** Selected transverse sections showing fibronectin deposition (green) and centrosome positioning (purple). Higher magnifications of the boxed regions are shown in A’. Scale bars = 20 μm. **(B)** Selected transverse sections showing Laminin deposition (red) and Golgi polarization (green). Higher magnifications of the boxed regions are shown in B’. Scale bars = 20 μm. **(C)** Selected transverse sections showing Fibronectin deposition (purple) and N-Cadherin polarization (green). Higher magnifications of the boxed regions are shown in C’. Scale bars = 20 μm. **(D)** Selected transverse sections showing Laminin deposition (red) and ZO-1 polarization (green). Higher magnifications of the boxed regions are shown in C’. Scale bars = 20 μm. **(E)** Selected transverse sections stained for the centrosomes (purple), N-cadherin (red) and aPKC (green) at the first steps of peripheral cell polarisation. The yellow arrow indicates apically located centrosomes with no apical protein accumulation. Scale bars = 10 μm. **(F)** Selected transverse sections stained for the centrosomes (purple), N-cadherin (red) and aPKC (green) at the first steps of peripheral cell polarisation. Yellow arrows indicate apically located centrosomes, with apical N-cadherin and/or aPKC. Scale bars = 10 μm. **(G)** Selected transverse sections stained for the centrosomes (purple), N-cadherin (red) and aPKC (green) at the first steps of peripheral cell polarisation. The yellow arrow indicates the apical position of centrosomes, with apical N-cadherin and aPKC protein accumulation. Scale bars = 10 μm. **(H)** Selected transverse sections stained for the centrosomes (purple), the basal protein β1 integrin (green) and the apical protein N-cadherin (red). β1 integrin is basally located in all cells with apically located centrosomes. The yellow arrow indicates apically located centrosomes, with apical N-cadherin and/or aPKC accumulation. Scale bars = 10 μm. **(I)** Plots the percentage of polarizing peripheral chord cells presenting only apical centrosome (FOP); apical centrosome and N-cadherin; apical centrosome, N-cadherin and aPKC; or apical centrosome, N-cadherin and ZO-1 (plot shows the mean±sem; n=100, 60, 87 cells from 10 embryos). **(J)** Plots the percentage of polarizing peripheral chord cells presenting apical centrosome (FOP) and basal β1 integrin or apical centrosome, basal β1 integrin and N-cadherin (plot shows the mean±sem; n=84 cells from 10 embryos). **(K-M)** Schemes showing the measurements performed in I,J. Cell nuclei are depicted in blue, centrosomes in purple (FOP) and the cis-Golgi in green (GM130).

**SUPPLEMENTARY FIGURE S5:**
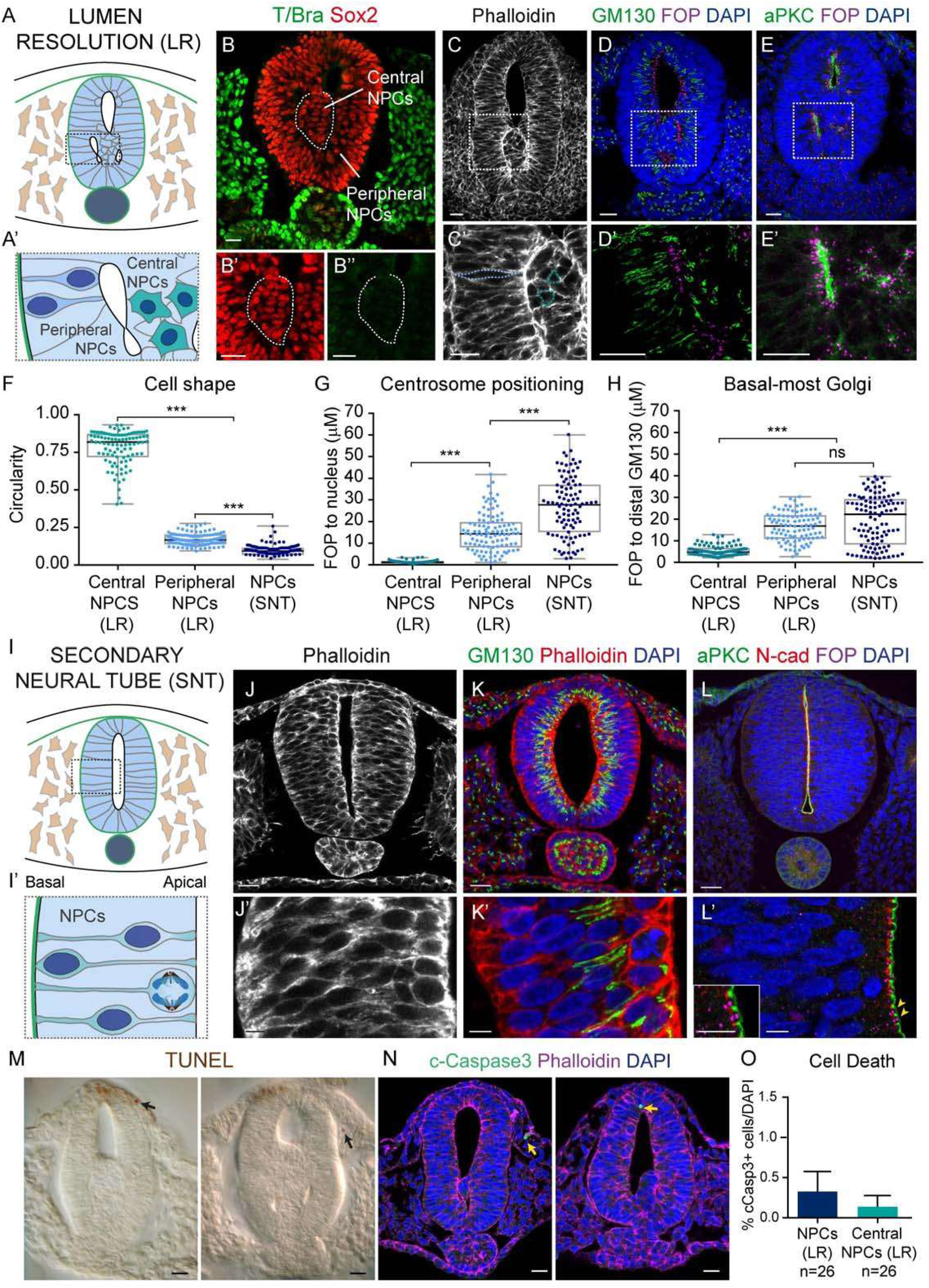
Central NPCs at the stage of lumen resolution maintain their mesenchymal characteristics. **(A)** Scheme of the cellular events occurring in secondary lumen resolution (LR). The basement membrane appears in green, and the two populations of peripheral and central NPCs are shown in A’. **(B)** Selected transverse sections in the LR phase stained for Sox2 (red) and T/Bra (green). Both central and peripheral NPCs are Sox2^+^ and T/Bra^-^ at this stage. The dotted white line defines the central NPCs and a higher magnification is shown in B’, B’’. Scale bars = 20 μm. **(C)** Selected transverse sections stained for the actin (white). In a higher magnification of the boxed region shown in C’, the dotted blue lines define the shape of peripheral and central NPCs. Scale bars = 20 μm. **(D)** Selected transverse sections stained for centrosomes (purple) and the cis-Golgi apparatus (green). DAPI (blue) stains the nuclei and a higher magnification of the boxed region is shown in D’. Scale bars = 20 μm. **(E)** Selected transverse sections stained for the centrosomes (purple) and the polarity protein aPKC (green). DAPI (blue) stains the nuclei and a higher magnification of the boxed region is shown E’. Scale bars = 20 μm. **(F)** Shape/circularity of central and peripheral NPCs at the LR stage, and of NPCs once the secondary neural tube (SNT) is fully formed (bold horizontal lines show the median; n=100, 100, 100 cells from 10 embryos): ***p<0.001 Kruskal-Wallis test. **(G)** The distance from the apical FOP to the nucleus in central and peripheral NPCs at the LR stage and in NPCs of the formed SNT (bold horizontal lines show the median; n=100, 100, 100 cells from 10 embryos): ***p<0.001 Kruskal-Wallis test. **(H)** The distance from FOP to the basal-most Golgi apparatus (distal GM130) in central and peripheral NPCs at the LR stage and in NPCs of the formed SNT (bold horizontal lines show the median; n=100, 100, 110 cells from 10 embryos): ***p<0.001 Kruskal-Wallis test. **(I)** Scheme of the formed SNT produced by SN with a single central lumen surrounded by dividing epithelial NPCs, as shown in I’. The apical and basal surfaces are indicated, and the basal membrane is green. **(J)** Selected transverse sections of the formed SNT stained for actin (white) and a higher magnification shown in J’. Dotted lines define the shape of central and peripheral NPCs. Scale bars = 20 μm (J), 5 μm (J’). **(K)** Selected transverse sections of the formed SNT stained for actin (red) and the cis-Golgi apparatus (green). DAPI (blue) stains the nuclei and a higher magnification is shown in K’. Scale bars = 20 μm (K), 5 μm (K’). **(L)** Selected transverse sections stained for centrosomes (purple), the polarity protein aPKC (green) and the junctional protein N-cadherin (red). DAPI (blue) stains the nuclei and a higher magnification is shown in L’, in which the yellow arrows indicate the two apical complexes shown at the bottom left. Scale bars = 20 μm (L), 5 μm (L’).

**SUPPLEMENTARY FIGURE S6:**
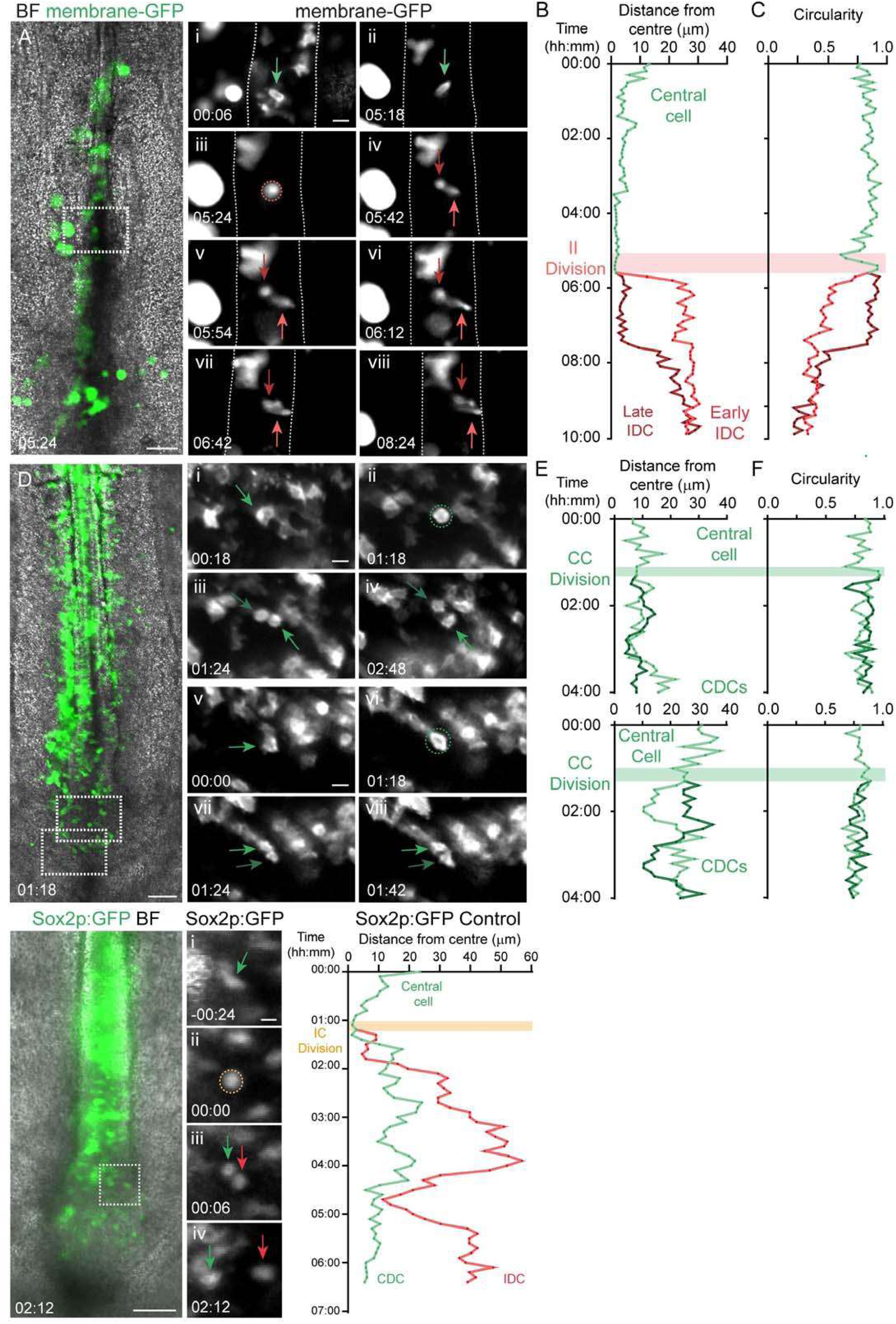
The majority of sh-SMAD3 divisions generate two central daughter cells that fail to intercalate into the lateral walls of the forming SNT. **(A)** Drawing of an electroporated HH15 chick embryo showing the cranio-caudal levels in B-E. **(B)** Selected frame of time-lapse movies of a Sox2p:GFP electroporated control embryo at the cranio-caudal level indicated in A. Red dotted lines defines the apico-basal surfaces of the SNT. Scale bar = 100 μm. **(C**,**D)** Selected frames of time-lapse movies of sh-SMAD3 Sox2p:GFP electroporated control embryos at the cranio-caudal levels indicated in A. The frames are the last time points of the mitotic sequences in G,H, as indicated by the boxed regions. Red dotted lines define the apico-basal surfaces of the SNT. Note the abnormal accumulation of cells in the centre of the SNT in C and D. Scale bars = 100 μm. **(E**,**F)** Selected frames of movies of sh-SMAD3 Sox2p:GFP showing three different cell divisions at the cranio-caudal positions indicated in C,D. The dotted circles indicate the mitosis (CC divisions, green), the green arrows point to the central cells. **(G**,**H)** Plots of the distance from the centre during the time-lapse movie of shSMAD3 Sox2p:GFP cells. All daughter cells remain close to the centre (both dark and light green). CDC, central daughter cell; IDC, intercalating daughter cell.

## SUPPLEMENTARY MOVIE LEGENDS

**Movie S1**. Transversal z-stack of the stage HH15 chick embryo in Figure 4E, advancing from the caudal to cranial sections. ZO-1 (green) shows the emerging lumen foci that coalesce into a single central cavity. Laminin (purple) stains the growing basal lamina and DAPI (blue) stains the cell nuclei. Related to Figure 4E.

**Movie S2**. 3D reconstruction of a stage HH15 chick embryo secondary lumen. The single cranial lumen caudally splits into one dorsal-central lumen and two ventral-lateral lumens. Related to Figure 4F.

**Movie S3**. In vivo time-lapse imaging of a membrane:GFP electroporated embryo showing a central cell (blue dot) that undergoes IC cell division and generates one intercalating daughter cell (red dot) and one daughter cell that remains in the centre (green dot). Related to Figure 5A,E.

**Movie S4**. In vivo time-lapse imaging of a membrane:GFP electroporated embryo showing a central cell (blue dot) that undergoes II cell division and generates two intercalating daughter cells (red and green dots). Related to Figure 5D and Sup Figure 6A-C.

**Movie S5**. In vivo time-lapse imaging of a membrane:GFP electroporated embryo showing various CC cell divisions. Related to Figures 5F and Sup Figure 6D-F.

**Movie S6**. In vivo time-lapse imaging of a pSox2:GFP control electroporated embryo showing a central cell (blue dot) that undergoes IC cell division and generates one intercalating daughter cell (red dot) and one daughter cell that remains in the centre (green dot). Related to Figure 32A-C.

**Movie S7**. In vivo time-lapse imaging of a sh-SMAD3 pSox2:GFP electroporated embryo where a central cell (blue dot) undergoes mitosis and generates two central daughter cells that finally elongate at a central position (red and green dots). Related to Figure 6C,F.

**Movie S8**. In vivo time-lapse imaging of a sh-SMAD3 pSox2:GFP electroporated embryo where a central cell (blue dot) undergoes mitosis and generates two central daughter cells that finally elongate at a central position (red and green dots). Related to Figure 6D,G.

## STAR ★ METHODS

### KEY RESOURCES TABLE

### LEAD CONTACT AND MATERIALS AVAILABILITY

Further information and requests for resources and reagents should be directed to and will be fulfilled by the Lead Contact, Elisa Martí.

### EXPERIMENTAL MODEL AND SUBJECT DETAILS

#### Chick Embryos

Fertilized eggs from the White-Leghorn strain of chickens were incubated horizontally at 38.5°C in an atmosphere of 70% humidity. Embryos were staged following morphological criteria (Hamburger and Hamilton, 1992).

#### Chick in ovo electroporation

An Intracel Dual Pulse (TSS-20 Ovodyne) electroporator equipped with a footswitch was used to generate electric pulses. We separated a pair of platinum commercial electrodes (CUY610P1.5-1, Nepagene) and only used one side as the positive electrode. We incorporated a sharpened and bent 90° tungsten needle (Fine Science Tools) into a holder and used it as the negative ‘microelectrode’

Eggs were horizontally incubated at 38.5°C in an atmosphere of 70% humidity until HH9 stage. DNA plasmids were diluted at 0.05-2 - g/ -1 in 60% sucrose in sigma H2O(Sigma Aldrich, W4502) with 50 ng/ml of Fast Green FCF (Sigma-Aldrich, F7258). Before manipulation, 5ml of albumen was removed from the egg with a syringe and a window was opened at the top of the shell to visualize the embryo. Thin forceps were used to open a small hole in the posterior region of the area opaca, just outside of the area pellucida 200ml of 1% Penicillin/Streptomycin (P/S) (Gibco, 15070063) were poured on top of the embryo to improve electrode conductivity. DNA solution was then injected onto the epiblast with a glass capillary by blowing air through an aspirator tube (Sigma-Aldrich, A5177-5EA). DNA was introduced with a glass capillary needle (GD-1, Narishige; made with Narishige PC-10 glass capillary puller) into the small concave region at their posterior end of the stage HH9 embryo, where the neural tube is still open. The platinum electrode connected to the positive lead (+) was carefully inserted below the embryo through the hole made previously, parallel to its antero-posterior axis. The tungsten microelectrode connected to the negative lead (-) was then positioned on top of the embryo, also in parallel to its antero-posterior axis. Five 50 ms square pulses of 5V at intervals of 50 ms were delivered. The window in the shell was finally sealed with plastic tape and embryos were incubated until the desired stage.

### METHOD DETAILS

#### Chick embryo culture mounting for in vivo imaging

Filter paper rings were prepared from 2 ⨯ 2 cm squares of Whatman grade 1 filter paper (Sigma-Aldrich, WHA1001325) in which a clover-leaf shaped hole was made in the centre with a paper punch, cutting the corners so that they fit in the round imaging plates. Imaging plates were also prepared in advanced by bedding several Millicell cell culture plate inserts (0.4 mm: Millipore, PICMORG50) with an Agar/Albumen mix.

The tape-sealed window in the egg was reopened and the thick albumen surrounding and covering the embryo carefully removed with a soft tissue. A paper ring was placed on top of the vitelline membrane so that the embryo located in the center of the clover-shaped hole, the vitelline membrane was cut through and around the whole perimeter of the filter paper ring and finally the filter with the embryo attached was pulled away from the yolk. Embryos with the best overall morphology and the greatest level of transgene expression were selected for imaging and transferred ventral side up to the imaging plates.

Imaging was performed inside a culture chamber created from a Corning ® Costar ® polystyrene 6-well plate (Sigma, CLS3736). To favour the optics, the plastic in the lid was replaced with glass. Each well of the culture chamber was filled with 1.5 mL of a solution of 5 ml thin albumen and 5 ml of 123 mM NaCl, the embryos in the imaging plates were transferred to the wells of the culture chamber and 1xPBS was added in between wells to maintain a moist environment inside the culture chamber. The culture chamber was finally sealed with electrical insulation tape.

#### In vivo time-lapse imaging

Embryos were visualised under an upright wide-field microscope Axio Imager 2 (Zeiss) equipped with a motorized stage and an incubation chamber. The temperature was set to 39.5 °C so that the temperature at the level of the embryo was around 37.5 °C. The Experiment designer module of version 2.3 blue edition of the ZEN software (Zeiss) (RRID: SCR_013672) was used to set up the acquisition. For 5x objective, 10 z images every 10 minutes for 100 loops were acquired with a resolution of 1024⨯1024 binning 4⨯4. For 20x objective, 10 z images every 6 minutes for 150 loops were acquired with a resolution of 1024⨯1024 binning 4⨯4. The images of each embryo acquired were first time-stitched with the ZEN software (Zeiss) and then exported to Image J/Fiji software for image processing and analysis.

#### Whole-mount immunohistochemistry

In toto embryo immunostaining procedure was carried out as follows: Chick embryos were removed from the egg at stage HH15 and fixed in 5ml 4% paraformaldehyde (PFA) (Sigma-Aldrich, 16005) in 1xPhosphate Buffered Saline (PBS) for 2 hours at room temperature (RT) or overnight at 4°C. Embryos were transferred to a 2ml tube, using a Pasteur pipette with the end cut off. Embryos were washed 3 ⨯30 min in 0.5% Triton-X-100 (Sigma-Aldrich, X100) in PBS (PBT).

Embryos were incubated in blocking solution consisting of 0.5% PBT + 1% Albumin from Bovine Serum (BSA, Sigma-Aldrich, 9048-46-8), 0.2% sodium azide (Sigma-Aldrich, S2002) for 1h at RT. Embryos were incubated in blocking solution with primary antibody for 2 to 3 days at 4°C with gentle shaking. Following incubation, embryos were washed 3 ⨯ 1h in 0.5% PBT. Embryos were then incubated in blocking solution with secondary antibodies for 2 days at 4°C with gentle shaking. After washing, embryos were incubated overnight at 4oC with DAPI (1:1000) (Sigma) in 0.5% PBT. Finally, embryos were initially washed 3 ⨯ 10 min in PBT, followed by 3 longer 30 min washes, transferred to PBS and stocked at 4°C. A full list of antibodies used in this study can be found in the Key Resources Table.

#### Free-floating sections immunohistochemistry

Immunostaining of transversal vibratome sections was carried out as follows: Chick embryos were removed from the egg at stage HH15 and fixed in 4% PFA in 1xPBS for 2 hours at RT or 4 h at 4°C. Embryos were embedded in plastic moulds with a warm 5% agarose - 10% sucrose matrix and cooled down to solidify. Agarose embryo-blocks were sectioned at 50-100µm thickness in a Leica Vibratome (VT1000S), obtaining free-floating transversal sections. Sections were washed 3 ⨯ 5 min in PBT (PBS + 0.1% Triton-X-100). Sections were incubated in blocking solution (10% BSA in PBT) for at least 30min at RT. Sections were incubated in antibody solution (1% BSA in PBT) with primary antibody overnight at 4°C with gentle shaking. Following incubation, sections were washed 3 ⨯ 10 min in PBT. Sections were then incubated in secondary antibodies in antibody solution for 2 hours at room temperature. Finally, embryos were initially washed 3 ⨯ 10 min in PBT washes and transferred to water for glass-slide mounting, and covered by Mowiol (Sigma-Aldrich, 81381) and a glass-coverslip.

Counter-stains were added during incubation with secondary antibody. DAPI (1:5000) was used to visualise nuclei (Sigma-Aldrich, D9542). TRITC conjugated phalloidin (1:1000) was used to visualize F-actin/tissue structure (Sigma-Aldrich, P1951). A full list of antibodies used in this study can be found in the <u>Key Resources Table</u>.

#### In situ hybridization

Embryos were removed from the egg at stage HH15 and fixed overnight at 4°C in 4% PFA diluted in 1xPBS. The next day embryos were dehydrated with a series of increasing methanol concentration solutions (25%, 50%, 75% and 100% methanol). Embryos were then stored at -20 °C for at least overnight. Whole-mount in situ hybridisation was performed following standard procedures with the InsituPro VSi robot (Intavis). Each condition was replicated in two wells with 3-4 embryos each. Probes from the chicken EST project (http://www.chick.manchester.ac.uk/) were used at 1:200. Sonic hedgehog probe was always used as positive control. Hybridized embryos were post-fixed in 4% PFA, rinsed in PBT and embedded in plastic moulds with a warm 5% agarose - 10% sucrose matrix and cooled down to solidify. Agarose embryo-blocks were sectioned at 50µm thickness in a Leica Vibratome, obtaining free-floating transversal sections. Finally, sections were transferred to water for glass-slide mounting, and covered by Mowiol and a glass-coverslip. A full list of probes used in this study can be found in the Key Resources Table.

#### TUNEL staining in free-floating sections

The deoxynucleotidyl transferase–mediated deoxyuridinetriphosphate nick end labelling (TUNEL) assay was used to detect programmed cell death by apoptosis. The TUNEL assay was performed using the In situ cell death detection kit POD (Roche, 11 684 817 910) following the manufacturer instructions with some modifications. Embryos were removed from the egg at stage HH15 and fixed overnight at 4°C in 4% PFA diluted in 1xPBS. The next day embryos were dehydrated with a series of solutions with increasing methanol concentration (25%, 50%, 75% and 100% methanol). Embryos were stored at - 20 °C for at least overnight and up to six months. Embryos were rehydrated and embedded in plastic moulds with a warm 5% agarose - 10% sucrose matrix and cooled down to solidify. Agarose embryo-blocks were sectioned at 50µm thickness in a Leica Vibratome, obtaining free-floating transversal sections. TUNEL staining was then performed and the most posterior sections were used as positive controls. Colour was developed using DAB substrate in a solution containing 0.3% H2O2, prepared following the manufacturer instructions (Sigma-Aldrich, 7411-49-6). DAB reaction was stopped by washing a few times in PBS pH=7. Finally, TUNEL stained sections were transferred to water for glass-slide mounting, and covered by Mowiol and a glass-coverslip.

### QUANTIFICATION AND STATISTICAL ANALYSIS

#### 3D lumen reconstruction

Raw whole-mount confocal data was exported to the Imaris software (Bitplane) (RRID:SCR_007370). The secondary forming lumen was reconstructed using the Contour Surface tool. The 3D structure was extracted by manually drawing the lumen contour, visible with ZO-1 immunostaining, on consecutive 2D z-slices.

Image analysis and quantifications

#### General Image Analysis

Raw confocal data was exported to ImageJ/FiJi (http://rsbweb.nih.gov/ij/)(RRID: SCR_003070) to be processed and analysed (Rueden et al., 2017; Schindelin et al., 2012). Projections of z-stacks are maximum projections unless otherwise indicated. Figures and schemes were generated using Adobe Illustrator CS5 (RRID: SCR_014199).

#### Quantifications in transversal sections

#### Nuclear SMAD3 intensity

Sh-SMAD3 and pSUPER control vectors were co-electroporated with H2B-RFP and stained with an antibody against endogenous SMAD3. Images from both conditions were acquired with the same laser and gain parameters. The polygon selection tool of ImageJ was used to delineate H2B-RFP+ cell nuclei and the integrated density was measured. Results are presented in GraphPad Prism 6 box & whisker plots.

#### Cell death

Cleaved-Caspase3 (c-Caspase3) antibody was used to detect apoptosis in fixed transversal sections of stage HH15 chick embryos. For WT quantifications, we counted both the number of c-Capase3+ cells and the number of total DAPI cells. For electroporated embryos, we counted c-Caspase3+ cells, H2B-RFP+ cells and total cells (DAPI). Percentages were then calculated and presented in GraphPad Prism 6 bar graphs.

#### Proliferation

Phospho-histone 3 (pH3) antibody was used to detect mitotic cells in fixed transversal sections of stage HH15 chick embryos. For WT quantifications, we counted both the number of pH3+ cells and the number of total DAPI cells. For electroporated embryos, we counted pH3+ cells, H2B-RFP+ cells and total cells (DAPI). Percentages were then calculated and presented in GraphPad Prism 6 bar graphs.

#### Sox2 and T/Bra nuclear intensities

Images from fixed transversal sections of stage HH15 chick embryos stained for Sox2 and T/Bra antibodies were acquired with the same gain and laser parameters. The area, integrated density and mean grey value were measured for each nucleus and for three neighbouring selections with no fluorescence (background measurements). The level of fluorescence in the nucleus was then determined with the corrected total cell fluorescence (CTCF). CTCF is calculated with the formula CTCF= Integrated nuclear density – (Area of selected nucleus x Mean fluorescence of background readings). Log10 (CTCF) was finally calculated and represented in GraphPad Prism 6 box & whisker plots.

Images from fixed stage HH15 transversal sections stained for Sox2, T/Bra and Fibronectin were used to correlate nuclear intensities (CTCF) with cell distance from the basement membrane (BM). A straight line was drawn with the ImageJ command from the centre of the nucleus to the closest Fibronectin staining and distance was measured. Results are presented in GraphPad Prism 6 linear regression plots. We considered peripheral cells those in contact with the BM and central cells those located further than 40μM from BM.

Sh-SMAD3 and H2B-RFP control vectors were electroporated and stained with an antibody against Sox2. Images were acquired with the same parameters and analysed. The polygon selection tool of ImageJ was used to delineate electroporated cell nuclei and the integrated density of Sox2 nuclear staining was measured. For each selected H2B-RFP+ positive nucleus, three nucleus of non-electroporated neighbouring cells (negative for H2B-RFP) were also delimited and their Sox2 integrated density measured. The following ratio was then calculated: integrated density of nuclear Sox2 H2B-RFP + cell/mean integrated density of three H2B-RFP-neighbouring cells. Results are presented in GraphPad Prism 6 box & whisker plots.

#### Cell shape

Actin staining (Phalloidin) was used to visualize cell shape in WT, pSUPER control and sh-SMAD3 transversal sections. Cells were delimited with the polygon selection tool and cell shape was quantified by measuring cell circularity, a parameter included in ImageJ Shape descriptors. Circularity is calculated with the formula 4π×[Area]/[Perimeter]2, with a value of 1.0 indicating a perfect circle. As the value approaches 0.0, it indicates an increasingly elongated shape. Results are presented in GraphPad Prism 6 box & whisker plots.

#### Centrosome positioning

The centrosomes were visualized with FOP antibody and DAPI was used to stain the nucleus in WT, pSUPER control and sh-SMAD3 transversal sections. The straight-line tool of ImageJ was used to draw a line from the centrosomes to the edge of the nucleus and the distance was measured. Results are presented in GraphPad Prism 6 box & whisker plots.

#### Golgi measurements; Basal-most Golgi

The centrosomes and the Golgi apparatus were visualized with FOP and GM130 antibodies, respectively, in HH15 WT transversal sections. The straight-line tool of ImageJ was used to draw a line from the centrosomes to the distal end of GM130 staining and the distance was measured. Results are presented in GraphPad Prism 6 box & whisker plots showing all points, median and interquartile range.

#### Golgi extension

The Golgi apparatus was visualized with GM130 antibody in pSUPER control and sh-SMAD3 transversal sections. The straight-line tool of ImageJ was used to draw a line from the apical to the basal limit of GM130 staining and its length was measured. Results are presented in GraphPad Prism 6 box & whisker plots.

#### Sequence of protein polarisation

Fixed transversal confocal images of FOP, N-cadherin, ZO-1, aPKC and β1 Integrin in the polarising medullary cord were used to define the sequence of epithelial polarity acquisition. Co-stainings were used to analyse the presence or absence of the mentioned components in each cell. We calculated the percentage of cells with: i. only the centrosome localised apically; ii. apically localised centrosome and apical N-cadherin; iii. apical centrosome, apical N-cadherin and apical ZO-1; iv. apical centrosome, apical N-cadherin and apical aPKC; v. apical centrosome and basal β1 integrin and vi. apical centrosome, basal β1 integrin and apical N-cadherin. Results are represented in GraphPad Prism 6 stacked bar graphs.

#### Cell length and distance from lumen foci to basement membrane

Actin staining (phalloidin) and DAPI were used to visualize cell shape and cell nuclei in transversal control sections at the lumen initiation stage. The straight-line tool of ImageJ was used to draw a line along the length of the cell and its distance was measured. aPKC and Laminin were used to visualise the small lumen foci and the basal lamina, respectively, in control and sh-SMAD3 transversal sections at the lumen initiation stage. A straight line was drawn from the lumen foci to the closest laminin staining with the ImageJ command. Results are presented in GraphPad Prism 6 box & whisker plots.

#### SMAD apico-basal intensity profiles

CEP152-GFP was electroporated in order to detect the two centrioles in chick neuroepithelial cells. CEP152-GFP electroporated embryos were co-stained with polyglutamilated tubulin, to visualize the primary cilium, and SMAD3, phSMAD2/3 and SMAD2 antibodies to study their localisation. Intensity profiles were generated with the Plot profile ImageJ command by drawing a straight line from the apical tip of the primary cilium to the basal daughter centriole. Intensity for the stained proteins was then measured along the drawn line.

#### Quantifications in time-lapse movies; Distance to last pair of somites

The distance to the last pair of somites was measured in the generated movies (membrane-GFP or sh-SMAD3 and pSUPER control co-electroporated with Sox2p:GFP). Cell divisions of mesenchymal cells in the centre of the tissue were spotted and the first time point where mitotic rounding was detected was analysed. A straight line was drawn from the mitosis to the level of the last pair of somites, visible in bright-field images, with the ImageJ straight-line tool. Distance was measured and presented in GraphPad Prism 6 box & whisker plots.

#### Distance from centre

A straight line was drawn in the centre of the neural tube by following the lumen in all movie time points. membrane-GFP+, sh-SMAD3 Sox2p:GFP + or pSUPER control Sox2p:GFP + mitosis were spotted and tracked back until the beginning of the movie. The distance from the drawn midline to the analysed cell and their daughter cells was measured in each time point with the ImageJ straight-line tool. Results along time are presented in GraphPad Prism 6 linear regression graphs.

#### Circularity

membrane-GFP+, sh-SMAD3 Sox2p:GFP + or pSUPER control Sox2p:GFP + cell divisions were spotted in time-lapse movies and tracked back until the beginning of the movie. Cells were delimited with the polygon selection tool of Image J and cell shape was quantified by measuring cell circularity in each time point. Circularity is calculated with the formula 4π×[Area]/[Perimeter]2, with a value of 1.0 indicating a perfect circle. As the value approaches 0.0, it indicates an increasingly elongated shape. Results along time are presented in GraphPad Prism 6 linear regression graphs.

### STATISTICAL ANALYSIS

Quantitative data is expressed as mean±sem/SD or as median±IQR. Statistical analysis was performed using the GraphPad Prism 6 (RRID: SCR_002798). Significance was assessed by performing the Mann-Whitney test when comparing two populations or the Kruskal-Wallis when comparing more than two. In this later case, Dunn’s multiple comparisons test was also run. In the few cases were data followed a normal distribution, assessed with the D’Agostino Pearson omnibus normality test, one-way ANOVA was performed. In this later case, Tukey’s multiple comparisons test was also run (*p<0.05, **p<0.01, and ***p<0.001).

## REFERENCES

Aaku-Saraste, E., Hellwig, A., & Huttner, W. B. (1996). Loss of occludin and functional tight junctions, but not ZO-1, during neural tube closure--remodeling of the neuroepithelium prior to neurogenesis. Developmental Biology, 180(2), 664–679. https://doi.org/10.1006/dbio.1996.0336

Abbott, A. (2011). Tissue-bank shortage: Brain child. In Nature (Vol. 478, Issue 7370, pp. 442–443). https://doi.org/10.1038/478442a

Afonso, C., & Henrique, D. (2006). PAR3 acts as a molecular organizer to define the apical domain of chick neuroepithelial cells. J Cell Sci, 119(Pt 20), 4293–4304. https://doi.org/10.1242/jcs.03170

Bedzhov, I., & Zernicka-Goetz, M. (2014). Self-organizing properties of mouse pluripotent cells initiate morphogenesis upon implantation. Cell, 156(5), 1032–1044. https://doi.org/10.1016/j.cell.2014.01.023

Benazeraf, B., Beaupeux, M., Tchernookov, M., Wallingford, A., Salisbury, T., Shirtz, A., Shirtz, A., Huss, D., Pourquie, O., Francois, P., & Lansford, R. (2017). Multi-scale quantification of tissue behavior during amniote embryo axis elongation. Development. https://doi.org/10.1242/dev.150557

Benazeraf, B., Francois, P., Baker, R. E., Denans, N., Little, C. D., & Pourquie, O. (2010). A random cell motility gradient downstream of FGF controls elongation of an amniote embryo. Nature, 466(7303), 248–252. https://doi.org/10.1038/nature09151

Bhowmick, N. A., Ghiassi, M., Bakin, A., Aakre, M., Lundquist, C. A., Engel, M. E., Arteaga, C. L., & Moses, H. L. (2001). Transforming growth factor-β1 mediates epithelial to mesenchymal transdifferentiation through a RhoA-dependent mechanism. Molecular Biology of the Cell. https://doi.org/10.1091/mbc.12.1.27

Brown, K. A., Pietenpol, J. A., & Moses, H. L. (2007). A tale of two proteins: differential roles and regulation of Smad2 and Smad3 in TGF-beta signaling. Journal of Cellular Biochemistry, 101(1), 9–33. https://doi.org/10.1002/jcb.21255

Bryant, D. M., Roignot, J., Datta, A., Overeem, A. W., Kim, M., Yu, W., Peng, X., Eastburn, D. J., Ewald, A. J., Werb, Z., & Mostov, K. E. (2014). A molecular switch for the orientation of epithelial cell polarization. Dev Cell, 31(2), 171–187. https://doi.org/10.1016/j.devcel.2014.08.027

Cambray, N., & Wilson, V. (2007). Two distinct sources for a population of maturing axial progenitors. Development (Cambridge, England), 134(15), 2829–2840. https://doi.org/10.1242/dev.02877

Catala, M., Teillet, M. A., & Le Douarin, N. M. (1995). Organization and development of the tail bud analyzed with the quail-chick chimaera system. Mech Dev, 51(1), 51–65. https://www.ncbi.nlm.nih.gov/pubmed/7669693

Chenn, A., Zhang, Y. A., Chang, B. T., & McConnell, S. K. (1998). Intrinsic polarity of mammalian neuroepithelial cells. Molecular and Cellular Neurosciences, 11(4), 183–193. https://doi.org/10.1006/mcne.1998.0680

Chi, Q., Yin, T., Gregersen, H., Deng, X., Fan, Y., Zhao, J., Liao, D., & Wang, G. (2014). Rear actomyosin contractility-driven directional cell migration in three-dimensional matrices: a mechano-chemical coupling mechanism. Journal of the Royal Society, Interface, 11(95), 20131072. https://doi.org/10.1098/rsif.2013.1072

Colas, J. F., & Schoenwolf, G. C. (2001). Towards a cellular and molecular understanding of neurulation. Dev Dyn, 221(2), 117–145. https://doi.org/10.1002/dvdy.1144

Criley, B. B. (1969). Analysis of embryonic sources and mechanims of development of posterior levels of chick neural tubes. J Morphol, 128(4), 465–501. https://doi.org/10.1002/jmor.1051280406

Dady, A., Havis, E., Escriou, V., Catala, M., & Duband, J. L. (2014). Junctional neurulation: a unique developmental program shaping a discrete region of the spinal cord highly susceptible to neural tube defects. J Neurosci, 34(39), 13208–13221. https://doi.org/10.1523/JNEUROSCI.1850-14.2014

Denker, E., Sehring, I. M., Dong, B., Audisso, J., Mathiesen, B., & Jiang, D. (2015). Regulation by a TGFbeta-ROCK-actomyosin axis secures a non-linear lumen expansion that is essential for tubulogenesis. Development, 142(9), 1639–1650. https://doi.org/10.1242/dev.117150

Diez del Corral, R., Breitkreuz, D. N., & Storey, K. G. (2002). Onset of neuronal differentiation is regulated by paraxial mesoderm and requires attenuation of FGF signalling. Development (Cambridge, England), 129(7), 1681–1691.

Edlund, S., Landström, M., Heldin, C. H., & Aspenström, P. (2002). Transforming growth factor-β-induced mobilization of actin cytoskeleton requires signaling by small GTPases Cdc42 and RhoA. Molecular Biology of the Cell. https://doi.org/10.1091/mbc.01-08-0398

Garriock, R. J., Chalamalasetty, R. B., Kennedy, M. W., Canizales, L. C., Lewandoski, M., & Yamaguchi, T. P. (2015). Lineage tracing of neuromesodermal progenitors reveals novel Wnt-dependent roles in trunk progenitor cell maintenance and differentiation. Development (Cambridge, England), 142(9), 1628–1638. https://doi.org/10.1242/dev.111922

Gonzalez-Gobartt, E; Allio, G; Bénazéraf, B; Martí, E. (2020). In vivo analysis of the Mesenchymal-to-Epithelial transition during chick secondary neurulation. In Methods in Molecular Biology.

Gouti, M., Delile, J., Stamataki, D., Wymeersch, F. J., Huang, Y., Kleinjung, J., Wilson, V., & Briscoe, J. (2017). A Gene Regulatory Network Balances Neural and Mesoderm Specification during Vertebrate Trunk Development. Dev Cell, 41(3), 243–261 e7. https://doi.org/10.1016/j.devcel.2017.04.002

Gouti, M., Tsakiridis, A., Wymeersch, F. J., Huang, Y., Kleinjung, J., Wilson, V., & Briscoe, J. (2014). In vitro generation of neuromesodermal progenitors reveals distinct roles for wnt signalling in the specification of spinal cord and paraxial mesoderm identity. PLoS Biol, 12(8), e1001937. https://doi.org/10.1371/journal.pbio.1001937

Greene, N. D., & Copp, A. J. (2014). Neural tube defects. Annu Rev Neurosci, 37, 221–242. https://doi.org/10.1146/annurev-neuro-062012-170354

Griffith, C. M., Wiley, M. J., & Sanders, E. J. (1992). The vertebrate tail bud: three germ layers from one tissue. Anatomy and Embryology, 185(2), 101–113. https://doi.org/10.1007/bf00185911

Hamburger, V., & Hamilton, H. L. (1951). A series of normal stages in the development of the chick embryo. Journal of Morphology. https://doi.org/10.1002/jmor.1050880104

Henrique, D., Abranches, E., Verrier, L., & Storey, K. G. (2015). Neuromesodermal progenitors and the making of the spinal cord. Development, 142(17), 2864–2875. https://doi.org/10.1242/dev.119768

Kölliker, A. (1884). Die embryonalen Keimblä tter und die Gewebe. Z. Wiss. Zool., 40, 179–213.

Kondoh, H., & Takemoto, T. (2012). Axial stem cells deriving both posterior neural and mesodermal tissues during gastrulation. Curr Opin Genet Dev, 22(4), 374–380. https://doi.org/10.1016/j.gde.2012.03.006

Le Douarin, N. M., Teillet, M. A., & Catala, M. (1998). Neurulation in amniote vertebrates: a novel view deduced from the use of quail-chick chimeras. Int J Dev Biol, 42(7), 909–916. https://www.ncbi.nlm.nih.gov/pubmed/9853821

Le Dreau, G, & Marti, E. (2012). Dorsal-ventral patterning of the neural tube: a tale of three signals. Dev Neurobiol, 72(12), 1471–1481. https://doi.org/10.1002/dneu.22015

Le Dreau, G, Saade, M., Gutierrez-Vallejo, I., & Marti, E. (2014). The strength of SMAD1/5 activity determines the mode of stem cell division in the developing spinal cord. J Cell Biol, 204(4), 591–605. https://doi.org/10.1083/jcb.201307031

Le Dreau, Gwenvael, Garcia-Campmany, L., Rabadan, M. A., Ferronha, T., Tozer, S., Briscoe, J., & Marti, E. (2012). Canonical BMP7 activity is required for the generation of discrete neuronal populations in the dorsal spinal cord. Development (Cambridge, England), 139(2), 259–268. https://doi.org/10.1242/dev.074948

Luxardi, G., Marchal, L., Thome, V., & Kodjabachian, L. (2010). Distinct Xenopus Nodal ligands sequentially induce mesendoderm and control gastrulation movements in parallel to the Wnt/PCP pathway. Development (Cambridge, England), 137(3), 417–426. https://doi.org/10.1242/dev.039735

Marthiens, V., & ffrench-Constant, C. (2009). Adherens junction domains are split by asymmetric division of embryonic neural stem cells. EMBO Rep, 10(5), 515–520. https://doi.org/10.1038/embor.2009.36

Martin-Belmonte, F., Yu, W., Rodriguez-Fraticelli, A. E., Ewald, A. J., Werb, Z., Alonso, M. A., & Mostov, K. (2008). Cell-polarity dynamics controls the mechanism of lumen formation in epithelial morphogenesis. Curr Biol, 18(7), 507–513. https://doi.org/10.1016/j.cub.2008.02.076

Martin, B. L., & Kimelman, D. (2012). Canonical Wnt signaling dynamically controls multiple stem cell fate decisions during vertebrate body formation. Developmental Cell, 22(1), 223–232. https://doi.org/10.1016/j.devcel.2011.11.001

McGrew, M. J., Sherman, A., Lillico, S. G., Ellard, F. M., Radcliffe, P. A., Gilhooley, H. J., Mitrophanous, K. A., Cambray, N., Wilson, V., & Sang, H. (2008). Localised axial progenitor cell populations in the avian tail bud are not committed to a posterior Hox identity. Development, 135(13), 2289–2299. https://doi.org/10.1242/dev.022020

Metzis, V., Steinhauser, S., Pakanavicius, E., Gouti, M., Stamataki, D., Ivanovitch, K., Watson, T., Rayon, T., Mousavy Gharavy, S. N., Lovell-Badge, R., Luscombe, N. M., & Briscoe, J. (2018). Nervous System Regionalization Entails Axial Allocation before Neural Differentiation. Cell. https://doi.org/10.1016/j.cell.2018.09.040

Miguez, D. G., Gil-Guinon, E., Pons, S., & Marti, E. (2013). Smad2 and Smad3 cooperate and antagonize simultaneously in vertebrate neurogenesis. J Cell Sci, 126(Pt 23), 5335–5343. https://doi.org/10.1242/jcs.130435

Morris, J. K., Rankin, J., Draper, E. S., Kurinczuk, J. J., Springett, A., Tucker, D., Wellesley, D., Wreyford, B., & Wald, N. J. (2016). Prevention of neural tube defects in the UK: a missed opportunity. Arch Dis Child, 101(7), 604–607. https://doi.org/10.1136/archdischild-2015-309226

Moustakas, A, Souchelnytskyi, S., & Heldin, C. H. (2001). Smad regulation in TGF-beta signal transduction. Journal of Cell Science, 114(Pt 24), 4359–4369.

Moustakas, Aristidis, & Heldin, C.-H. (2002). From mono-to oligo-Smads: the heart of the matter in TGF-beta signal transduction. Genes & Development, 16(15), 1867–1871. https://doi.org/10.1101/gad.1016802

Nakamura, N., Rabouille, C., Watson, R., Nilsson, T., Hui, N., Slusarewicz, P., Kreis, T. E., & Warren, G. (1995). Characterization of a cis-Golgi matrix protein, GM130. The Journal of Cell Biology, 131(6 Pt 2), 1715–1726. https://doi.org/10.1083/jcb.131.6.1715

Nievelstein, R. A., Hartwig, N. G., Vermeij-Keers, C., & Valk, J. (1993). Embryonic development of the mammalian caudal neural tube. Teratology, 48(1), 21–31. https://doi.org/10.1002/tera.1420480106

Nowotschin, S., Ferrer-Vaquer, A., Concepcion, D., Papaioannou, V. E., & Hadjantonakis, A. K. (2012). Interaction of Wnt3a, Msgn1 and Tbx6 in neural versus paraxial mesoderm lineage commitment and paraxial mesoderm differentiation in the mouse embryo. Dev Biol, 367(1), 1–14. https://doi.org/10.1016/j.ydbio.2012.04.012

O’Rahilly, R., & Muller, F. (1994). Neurulation in the normal human embryo. Ciba Found Symp, 181, 70–82; discussion 82-9.

O’Rahilly, R., & Muller, F. (2002). The two sites of fusion of the neural folds and the two neuropores in the human embryo. Teratology, 65(4), 162–170. https://doi.org/10.1002/tera.10007

Olivera-Martinez, I., Harada, H., Halley, P. A., & Storey, K. G. (2012). Loss of FGF-dependent mesoderm identity and rise of endogenous retinoid signalling determine cessation of body axis elongation. PLoS Biol, 10(10), e1001415. https://doi.org/10.1371/journal.pbio.1001415

Ridley, A. J., Schwartz, M. A., Burridge, K., Firtel, R. A., Ginsberg, M. H., Borisy, G., Parsons, J. T., & Horwitz, A. R. (2003). Cell migration: integrating signals from front to back. Science (New York, N.Y.), 302(5651), 1704–1709. https://doi.org/10.1126/science.1092053

Rupp, P. A., Rongish, B. J., Czirok, A., & Little, C. D. (2003). Culturing of avian embryos for time-lapse imaging. Biotechniques, 34(2), 274–278. https://doi.org/10.2144/03342st01

Saade, M., Gutierrez-Vallejo, I., Le Dreau, G., Rabadan, M. A., Miguez, D. G., Buceta, J., & Marti, E. (2013). Sonic hedgehog signaling switches the mode of division in the developing nervous system. Cell Rep, 4(3), 492–503. https://doi.org/10.1016/j.celrep.2013.06.038

Saitsu, H., & Shiota, K. (2008). Involvement of the axially condensed tail bud mesenchyme in normal and abnormal human posterior neural tube development. Congenit Anom (Kyoto), 48(1), 1–6. https://doi.org/10.1111/j.1741-4520.2007.00178.x

Saitsu, H., Yamada, S., Uwabe, C., Ishibashi, M., & Shiota, K. (2004). Development of the posterior neural tube in human embryos. Anat Embryol (Berl), 209(2), 107–117. https://doi.org/10.1007/s00429-004-0421-2

Schoenwolf, G. C. (1984). Histological and ultrastructural studies of secondary neurulation in mouse embryos. Am J Anat, 169(4), 361–376. https://doi.org/10.1002/aja.1001690402

Schoenwolf, G. C., & Delongo, J. (1980). Ultrastructure of secondary neurulation in the chick embryo. Am J Anat, 158(1), 43–63. https://doi.org/10.1002/aja.1001580106

Schoenwolf, G. C., & Kelley, R. O. (1980). Characterization of intercellular junctions in the caudal portion of the developing neural tube of the chick embryo. Am J Anat, 158(1), 29–41. https://doi.org/10.1002/aja.1001580105

Shen, X., Li, J., Hu, P. P. C., Waddell, D., Zhang, J., & Wang, X. F. (2001). The Activity of Guanine Exchange Factor NET1 Is Essential for Transforming Growth Factor-β-mediated Stress Fiber Formation. Journal of Biological Chemistry. https://doi.org/10.1074/jbc.M009534200

Shi, Y., & Massagué, J. (2003). Mechanisms of TGF-beta signaling from cell membrane to the nucleus. Cell, 113(6), 685–700. https://doi.org/10.1016/s0092-8674(03)00432-x

Shimokita, E., & Takahashi, Y. (2011). Secondary neurulation: Fate-mapping and gene manipulation of the neural tube in tail bud. Dev Growth Differ, 53(3), 401–410. https://doi.org/10.1111/j.1440-169X.2011.01260.x

Shum, A. S., Tang, L. S., Copp, A. J., & Roelink, H. (2010). Lack of motor neuron differentiation is an intrinsic property of the mouse secondary neural tube. Dev Dyn, 239(12), 3192–3203. https://doi.org/10.1002/dvdy.22457

Takemoto, T., Uchikawa, M., Kamachi, Y., & Kondoh, H. (2006). Convergence of Wnt and FGF signals in the genesis of posterior neural plate through activation of the Sox2 enhancer N-1. Development (Cambridge, England), 133(2), 297–306. https://doi.org/10.1242/dev.02196

Taverna, E., Mora-Bermudez, F., Strzyz, P. J., Florio, M., Icha, J., Haffner, C., Norden, C., Wilsch-Brauninger, M., & Huttner, W. B. (2016). Non-canonical features of the Golgi apparatus in bipolar epithelial neural stem cells. Sci Rep, 6, 21206. https://doi.org/10.1038/srep21206

Theveneau, E., & Mayor, R. (2012). Neural crest delamination and migration: from epithelium-to-mesenchyme transition to collective cell migration. Developmental Biology, 366(1), 34–54. https://doi.org/10.1016/j.ydbio.2011.12.041

Tsakiridis, A., & Wilson, V. (2015). Assessing the bipotency of in vitro-derived neuromesodermal progenitors. F1000Research, 4, 100. https://doi.org/10.12688/f1000research.6345.2

Turner, D. A., Hayward, P. C., Baillie-Johnson, P., Rue, P., Broome, R., Faunes, F., & Martinez Arias, A. (2014). Wnt/beta-catenin and FGF signalling direct the specification and maintenance of a neuromesodermal axial progenitor in ensembles of mouse embryonic stem cells. Development, 141(22), 4243–4253. https://doi.org/10.1242/dev.112979

Tzouanacou, E., Wegener, A., Wymeersch, F. J., Wilson, V., & Nicolas, J. F. (2009). Redefining the progression of lineage segregations during mammalian embryogenesis by clonal analysis. Dev Cell, 17(3), 365–376. https://doi.org/10.1016/j.devcel.2009.08.002

Uchikawa, M., Ishida, Y., Takemoto, T., Kamachi, Y., & Kondoh, H. (2003). Functional analysis of chicken Sox2 enhancers highlights an array of diverse regulatory elements that are conserved in mammals. Dev Cell, 4(4), 509–519. https://www.ncbi.nlm.nih.gov/pubmed/12689590

Ulloa, F., & Briscoe, J. (2007). Morphogens and the control of cell proliferation and patterning in the spinal cord. Cell Cycle, 6(21), 2640–2649. https://doi.org/10.4161/cc.6.21.4822

Wymeersch, F. J., Huang, Y., Blin, G., Cambray, N., Wilkie, R., Wong, F. C. K., & Wilson, V. (2016). Position-dependent plasticity of distinct progenitor types in the primitive streak. ELife, 5, e10042. https://doi.org/10.7554/eLife.10042

Yamaguchi, T. P., Bradley, A., McMahon, A. P., & Jones, S. (1999). A Wnt5a pathway underlies outgrowth of multiple structures in the vertebrate embryo. Development (Cambridge, England), 126(6), 1211–1223.

Yan, X., Habedanck, R., & Nigg, E. A. (2006). A complex of two centrosomal proteins, CAP350 and FOP, cooperates with EB1 in microtubule anchoring. Molecular Biology of the Cell, 17(2), 634–644. https://doi.org/10.1091/mbc.e05-08-0810

Yang, J., & Weinberg, R. A. (2008). Epithelial-mesenchymal transition: at the crossroads of development and tumor metastasis. Dev Cell, 14(6), 818–829. https://doi.org/10.1016/j.devcel.2008.05.009

Yoshikawa, Y., Fujimori, T., McMahon, A. P., & Takada, S. (1997). Evidence That Absence ofWnt-3aSignaling Promotes Neuralization Instead of Paraxial Mesoderm Development in the Mouse. Developmental Biology, 183(2), 234–242. https://doi.org/10.1006/DBIO.1997.8502

